# Data-derived agents reveal dynamical reservoirs in mouse cortex for adaptive behavior

**DOI:** 10.64898/2026.03.03.709365

**Authors:** Siyan Zhou, Ryan P. Badman, Charlotte Arlt, Kanaka Rajan, Christopher D. Harvey

## Abstract

Animals generate behaviors that are robust to perturbations yet adaptable to changing conditions. How neural population dynamics support this balance between robustness and flexibility remains unclear. We address this question in goal-directed navigation by combining large-scale calcium imaging from mouse cortex with a data-derived modeling framework. We trained agents to navigate in a simulative environment while recapitulating mouse neural and behavioral data trial-by-trial. Data-derived agents discovered novel dynamics of chaotic attractors, characterized by intrinsically variable trajectories confined within overall goal-specific attracting landscapes. These dynamics support reliable goal achievement while maintaining a structured distribution of navigational trajectories. Circuit-level analyses and perturbations reveal mechanisms that stabilize chaos and enhance behavioral adaptability in the data-derived agents. Thus, through our new modeling approach that emphasizes closed-loop interactions between behavior and neural dynamics, we reveal chaos as a functional principle for flexible behavior.

## Introduction

Brains generate behaviors that are robust to perturbations yet adaptable to changing conditions. Even for behavioral outcomes that are stable and robust, the activity in populations of neurons often exhibits substantial variability across repeated trials and conditions ^1–7^. An interesting proposal is that the trial-to-trial variability in neural activity may play a functional role in flexible behavior, rather than reflecting noise that is simply averaged away.

Existing theories of neural dynamics typically emphasize either robustness or flexibility, but not both simultaneously. Classical attractor-based frameworks achieve reliable outcomes by contracting neural trajectories toward fixed points or low-dimensional manifolds, a mechanism that necessarily suppresses within-condition variability ^8–15^. Conversely, models that generate flexible or diverse trajectories often rely on transient or feedforward dynamics which are not designed to guarantee robustness to perturbations ^16–20^. As a result, current theories lack a dynamical regime that can simultaneously support stable behavioral outcomes and rich neural and behavioral variability under the closed-loop demands that animals naturally operate under.

Neural activity unfolds in continuous interaction with the environment, such that variability in neural dynamics shapes future sensory input, which in turn influences subsequent neural activity ^21–24^. Yet many widely used models of neural dynamics are primarily evaluated in open-loop settings, decoupled from the behavioral consequences of neural activity ^25,26^. Severing this interaction obscures how neural dynamics support behavior over extended trajectories and might systematically mischaracterize the functional role of variability.

Goal-directed navigation naturally exposes this challenge. Animals reliably reach spatial goals while expressing diverse trajectories across trials, even under similar environmental conditions ^1,2,27,28^. This dissociation between outcome reliability and path variability provides a clear setting in which to investigate how neural population dynamics can support stable behavioral goals without committing to fixed trajectories.

Here we show that chaotic neural dynamics provide a regime in which population activity supports reliable goal attainment while simultaneously generating diverse and adaptable trajectories. In closed-loop interaction, chaos enables structured exploration without compromising goal stability, rather than producing unstructured instability. This regime reconciles robustness and flexibility within a single set of neural dynamics.

Identifying this functional role of chaos requires models that are evaluated under closed-loop interaction with the environment. We achieved this by reconstructing the neural dynamics jointly with the closed-loop interactive behavior of navigation. In open-loop analyses that primarily evaluate trajectory predictability or stability, chaotic regimes can appear unstable or uninformative and are therefore often disfavored during model fitting or selection ^15,29,30^. Closed-loop evaluation reveals that these same regimes can support reliable behavior while preserving structured variability, a distinction that cannot be recovered by fitting neural activity alone.

Together, our results establish chaos as a functional principle of neural population dynamics supporting flexible behavior. More broadly, they suggest that balancing stability and adaptability in closed-loop tasks may rely on dynamical regimes that are systematically overlooked by standard analytical approaches.

## Results

### Closed-loop modeling of neural dynamics and behaviors

In sequential behavior tasks like navigation, animals can achieve their goals through a variety of behavioral sequences. This is permitted by these tasks by definition, as they admit or even encourage variable behavioral sequences that fulfill a higher-level goal and comply with certain task constraints (Fig. 1a). In navigation, the higher-level goal is to reach a destination location, and the task constraints manifest as the organization of the maze walls, as well as coupling laws of how locomotion actions move the animal’s location in the environment and induce different future state distributions and requirements on subsequent actions. These constraints are embedded in the closed-loop interaction between the nervous system and the environment. Examining the neural activity alone without acknowledging the closed-loop interaction may obscure how neural dynamics support the temporally extended behavior.

**Fig. 1.**
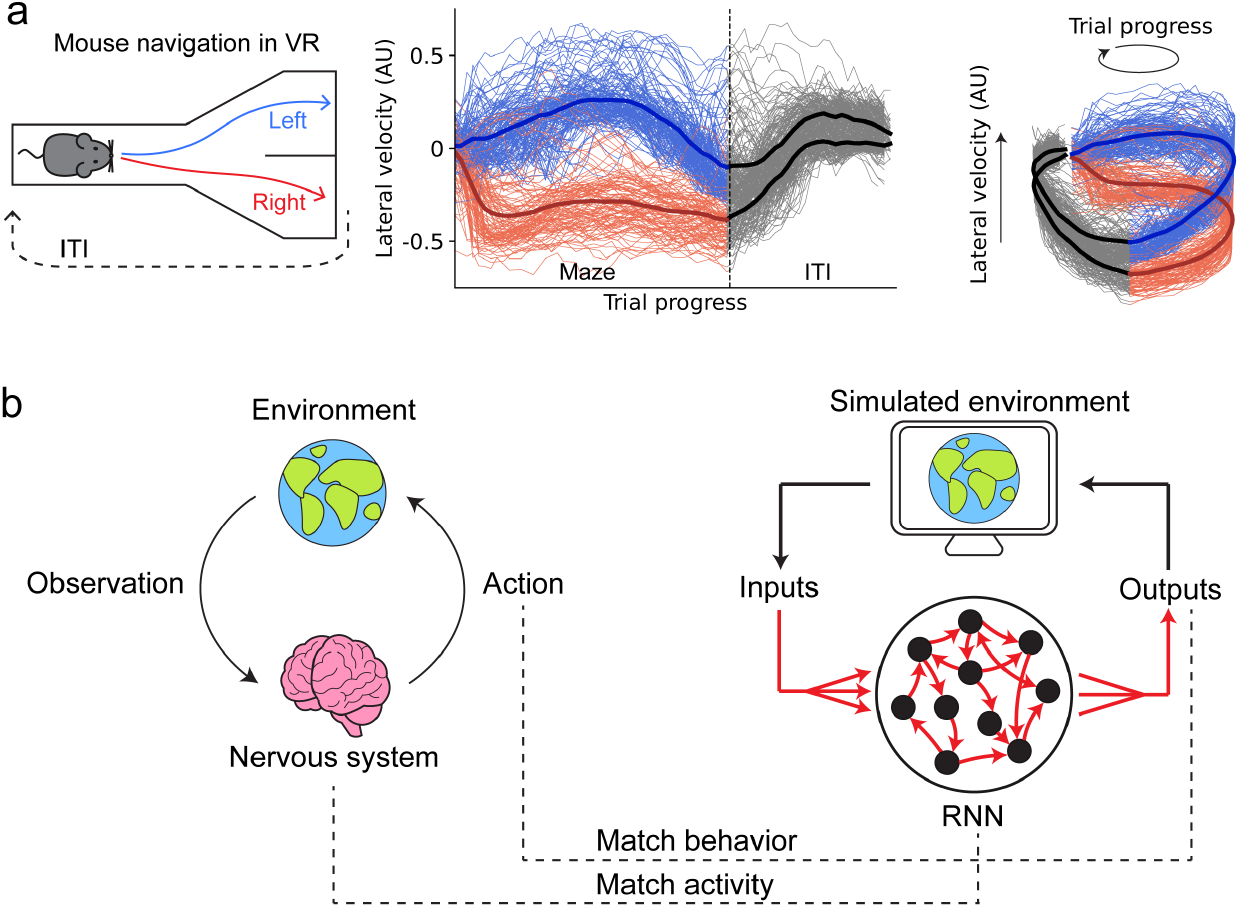
Data-derived agents jointly model neural dynamics and closed-loop behavior. **(a)** Left and middle: Mouse navigation in a virtual reality (VR) Y-maze shows diverse locomotor trajectories toward the same discrete choice. Thin lines indicate individual left or right trials in one mouse session; thick lines indicate trial-averaged trajectories. ITI, inter-trial interval. Right: Representing trial progress in polar coordinates reveals cycle-like geometry, but trajectories do not converge to fixed limit cycles, instead forming choice-specific distributions. **(b)** Schematic of ARCTIC (Activity Reconstruction in Closed loop). RNN-based agents are trained to perform cognitive tasks such as navigation in closed-loop simulated environments, while constrained by neural and behavioral data from real task-performing animals. Red arrows denote learnable connection weights.

To jointly model the underlying neural dynamics and the closed-loop interactive behavior, we developed a method that combines two commonly used modeling directions, neural dynamical systems reconstruction through data fitting ^19,31^, and agent-based modeling ^32^. Data-fitting models train dynamical systems like recurrent neural networks (RNN) to reconstruct experimentally recorded neural activity sequences. However, most existing approaches treat sensory inputs as independent processes, either pre-designed and fixed or co-optimized with the RNN weights as free parameters ^19,33,34^. This assumption of independence neglects the closed-loop environment interaction in navigation. On the other hand, task-performing agents are trained (usually by reinforcement learning) to perform tasks by acting in an environment and receiving sensory inputs according to their own actions in a closed loop ^32,35–40^. Yet these agents often aim to identify effective policies rather than reveal specific neural dynamics mechanisms. A common current practice is to compare internal representations of the agents with real animals in a *post hoc* manner, rather than using real neural dynamics to drive the agent’s behavior ^41^.

We propose ARCTIC (Activity Reconstruction in Closed loop) that combines data fitting with closed-loop agents (Fig. 1b). The approach features an environment-interacting agent that is constrained by both the neural activity and behavioral outputs recorded from real task-performing animals. There are two key technical advancements. First, in the spirit of the closed-loop environment interaction, neural activity reconstruction and behavior output optimization are trained simultaneously in an online manner ^42^. This helps to address error accumulation in the closed-loop setup. Second, the agent is trained on the neural and behavioral data of individual animal trials rather than averaged data. This enables the agent to recapitulate the distribution of possible sequences and the underlying neural mechanism for such variability (see Methods).

### Mice exhibit variable trajectories to the same choice in navigation

We studied mouse navigation to understand how a distribution of trajectories arises from neural dynamics. Mice were trained to perform a navigation decision-making task in a virtual reality Y-maze ^2,43^ (Fig. 2a). On each trial, one of two visual cues was presented in the Y-stem, and mice learned that each cue was associated with either a left or right turn at the Y-intersection to receive a reward. As mice performed the task, cellular-resolution calcium imaging via a large field-of-view, random-access microscope ^44^ was used to measure the activity of thousands of individual neurons across four cortical areas: primary visual cortex, posterior parietal cortex, retrosplenial cortex, and secondary motor cortex (Fig. 2b, Extended Data Fig. 1a). Mice performed the task with high accuracy across sessions (Fig. 2c). A portion of these experimental data was reported previously in ref. ^43^.

**Fig. 2.**
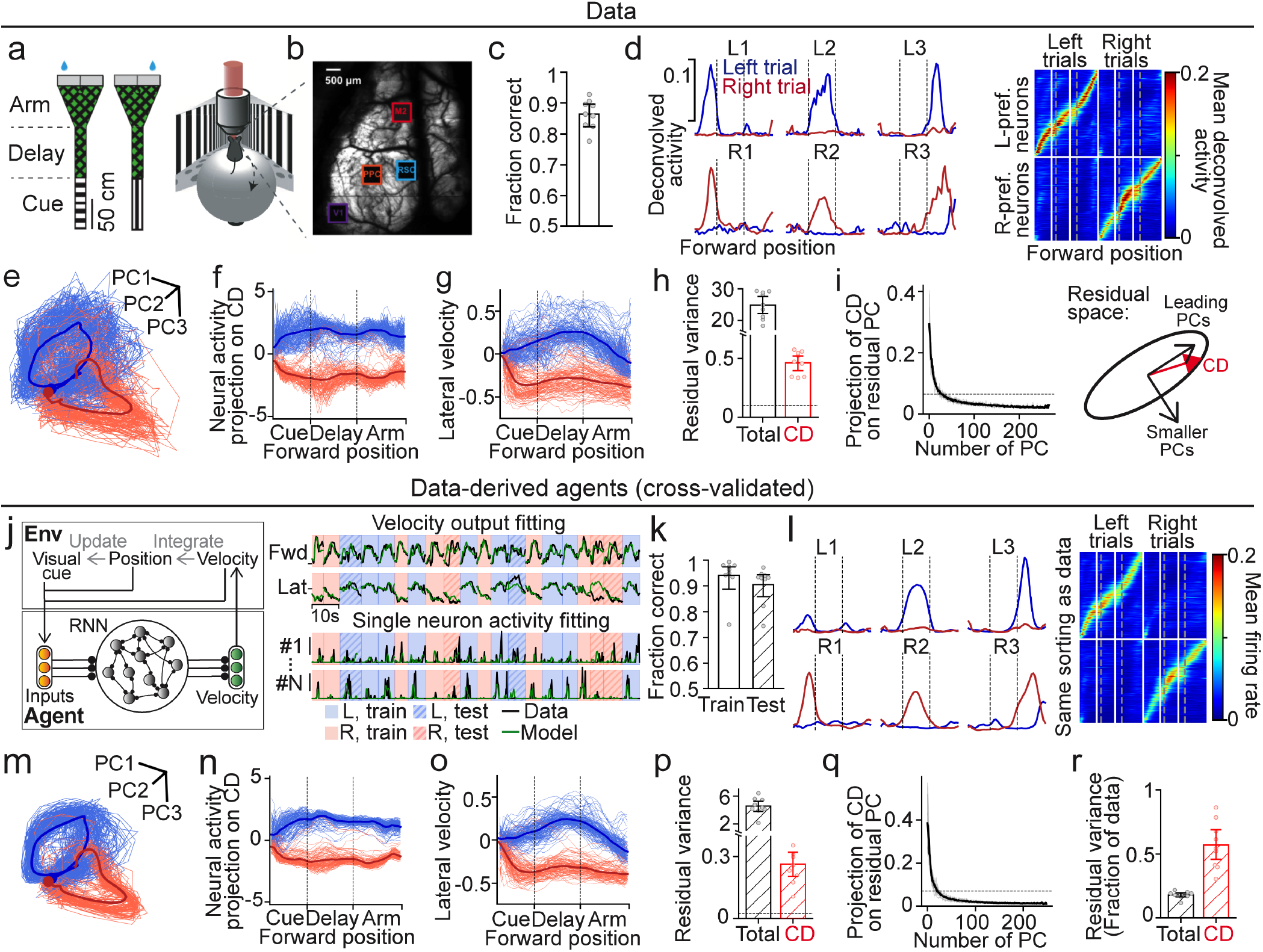
Data-derived agents recapitulate neural and behavioral trajectories during navigation. **(a)** Virtual reality Y-maze task. One of two visual cues is presented in the Y-stem, each associated with a left or right reward location. **(b)** Overview image showing the four dorsal cortical areas imaged simultaneously: V1, PPC, RSC, M2. **(c)** Mice achieved good behavioral performance across sessions (n = 2 mice, 9 sessions). **(d)** Left: Mean activity in left and right trials for three left- and three right-preferring example neurons. Dotted lines indicate boundaries between cue, delay, and arm epochs. Right: Mean activity of all choice-selective neurons pooled across areas and sessions, sorted by peak forward position in the preferred choice. **(e)** PCA embedding of individual correct left and right trials in an example session. Thick lines indicate trial means; dots mark starting points. **(f)** Neural activity projected on the choice dimension (CD) for the same session as in (e). **(g)** Lateral running velocity for individual trials in the same session. **(h)** Total residual variance of neural activity and residual variance projected on the CD. Each dot represents one session. Dotted line indicates chance-level CD projection averaged across sessions. **(i)** Projection of the CD on principal components (PCs) of the residual space. Black curve shows session average. Grey curves show individual sessions. Dotted line indicates chance-level projection averaged across sessions. Right: Illustration of CD alignment with leading residual PCs. **(j)** Left: Schematic of the environment-interacting RNN agent. Ball-headed arrows denote learnable weights. Right: The agent’s velocity output and single-neuron activity are fit to single-trial data. One agent is trained per crossvalidation fold for each session. **(k)** Agent performance on training and test trials (fraction correct). Each dot represents one session, averaged across five-fold cross-validation. In the following panels (l-r), the agents are evaluated for their neural activity and behavior in cross-validated test trials. **(l)** Left: Agentpredicted trial-mean activity for the six example neurons in (d). Right: Trial-mean activity of units pooled across agents, indexed as in (d). **(m-n)** Single-trial agent activity from the same session as in (e) projected on the PC space of the data (m) or the CD of the data (n). **(o)** Agent-generated lateral velocity in the same session. **(p)** Total residual variance generated by agents and its projection on the CD. Dotted line indicates chance level. **(q)** Projection of the CD on residual PCs of the agents. **(r)** Fraction of total residual variance in the data captured by the agents, compared with the fraction of residual variance on CD captured by the agents. Error bars in this and the following figures indicate 95% confidence interval obtained by bootstrapping.

Neural activity unfolded as choice-selective sequences ^45^ (Fig. 2d). Individual neurons were transiently active at specific points in each trial of the task. Different sequences of neurons were active on trials associated with different behavioral choices. Although population activity followed distinct trajectories that were separable for different choices, it exhibited substantial variability across trials of the same choice. This variability was concentrated along task-relevant dimensions of neural activity. It was evident in both the leading principal components of population activity and the choice dimension, defined as the axis that best separates neural activity for left versus right choices (Fig. 2e, f). Consistent with the neural variability, mice showed behavioral differences in left-right (lateral) running velocity across trials, even when navigating to the same goal location (Fig. 2g). This behavioral variability emerged because, by nature of the navigation task, the trial evolution is fully determined by the mouse. With the long running distance (2 m) and trial duration (5-10 s, Extended Data Fig. 1b), it is sensible that mice explored different trajectories and made variable movements in their running at different points across trials.

To isolate trial-to-trial neural variability, we subtracted the mean activity at each forward position in the maze from each individual trial, computed separately for left and right choice trials. The resulting residual activity reflects variability across repeated trials that is not explained by choice or forward position within the maze. If this trial-to-trial variance is unstructured, its projection onto the choice dimension would be small and near chance level. Instead, substantial residual variance, and its leading principal components, were strongly aligned with the choice dimension (Fig. 2h-i). This finding is consistent with previous work showing an alignment of signal and noise correlations in neural populations ^4,5,46^. Together, these results indicate that the trial-to-trial variability is task-relevant and unlikely to arise from unstructured noise or encoding of irrelevant stimuli or behaviors.

### Data-derived agents recapitulate variable trajectories without added noise

To investigate the mechanisms underlying these activity patterns, we applied ARCTIC to train data-derived navigation agents. Each agent has a densely connected RNN in which each unit corresponds to a single neuron recorded during a calcium imaging and behavioral session. Each unit was trained to reproduce the activity of a corresponding neuron ^19,29,31^, while the network as a whole was optimized to match the mouse’s forward and lateral running velocities (Fig. 2j). We trained one agent for each animal session. The agent interacted with a simulative environment analogously to how a mouse interacts with its environment. The agent received inputs from the environment corresponding to the visual cue and its current position in the maze. The agent’s velocity outputs then updated its position, thereby determining the next set of inputs. For each trial, the activity of units in the RNN and the position in the environment were initialized to the first empirical datapoint of that trial. The trained agent then autonomously generated the full neural and behavioral sequence of the trial, meaning its outputs arose entirely from internal dynamics and closed-loop interaction with the environment. After training, the agents performed the task with high accuracy (Fig. 2k).

The agents produced single-neuron activity and behavioral outputs that recapitulated key features of the mouse data. Strikingly, this was the case even in test trials held out from training. First, the agents generated distinct neural sequences and running trajectories for each trial type, matching the experimentally observed separation by mouse choice (Fig. 2l–o). Second, the agents recapitulated trial-to-trial variability in both neural activity and behavior. This variation emerged even within a single trial type, despite the absence of added noise (Fig. 2m–o; Extended Data Fig. 1f). Like the experimental data, the residual variance, which was computed after subtracting trial-averaged activity, was well aligned with the choice dimension (Fig. 2p–q). Also, the agents captured a larger fraction of the residual variance along the choice dimension compared to the overall residual variance, potentially because they ignored task-irrelevant components (Fig. 2r). Together, these results indicate that the data-derived agents developed intrinsic mechanisms for generating task-relevant trial-to-trial variability that was distinct from random noise, while maintaining high decision-making accuracy.

### Low-dimensional chaotic attractors

To probe a data-derived agent’s intrinsic dynamics, we perturbed its RNN activity along the choice dimension and tracked the evolution of the agent’s average response over time. Small perturbations decayed rapidly and preserved behavioral accuracy, whereas larger perturbations caused the activity to jump to the opposite trial type, resulting in incorrect choices (Fig. 3a; Extended Data Fig. 2a). To quantify the average convergence of neural activity following a perturbation, we computed the norm deviation of trial-averaged neural activity (NDM), defined as the squared difference between perturbed and unperturbed means across trials of the same choice. NDM decreased rapidly after a perturbation in correct trials, indicating that mean neural activity returned toward its original trajectory (Fig. 3b). As the perturbation amplitude increased, choice accuracy decreased nonlinearly, suggesting a boundary separating neural activity for the two choices (Extended Data Fig. 2b). Perturbations had a greater effect during the delay period than during the cue period, consistent with the idea that visual cues constrain neural activity early in the trial (Extended Data Fig. 2a). Together, these findings support the existence of separate attractor states for left and right choices. This is consistent with previous theories that attractor dynamics support reliable decision making ^8–12^.

**Fig. 3.**
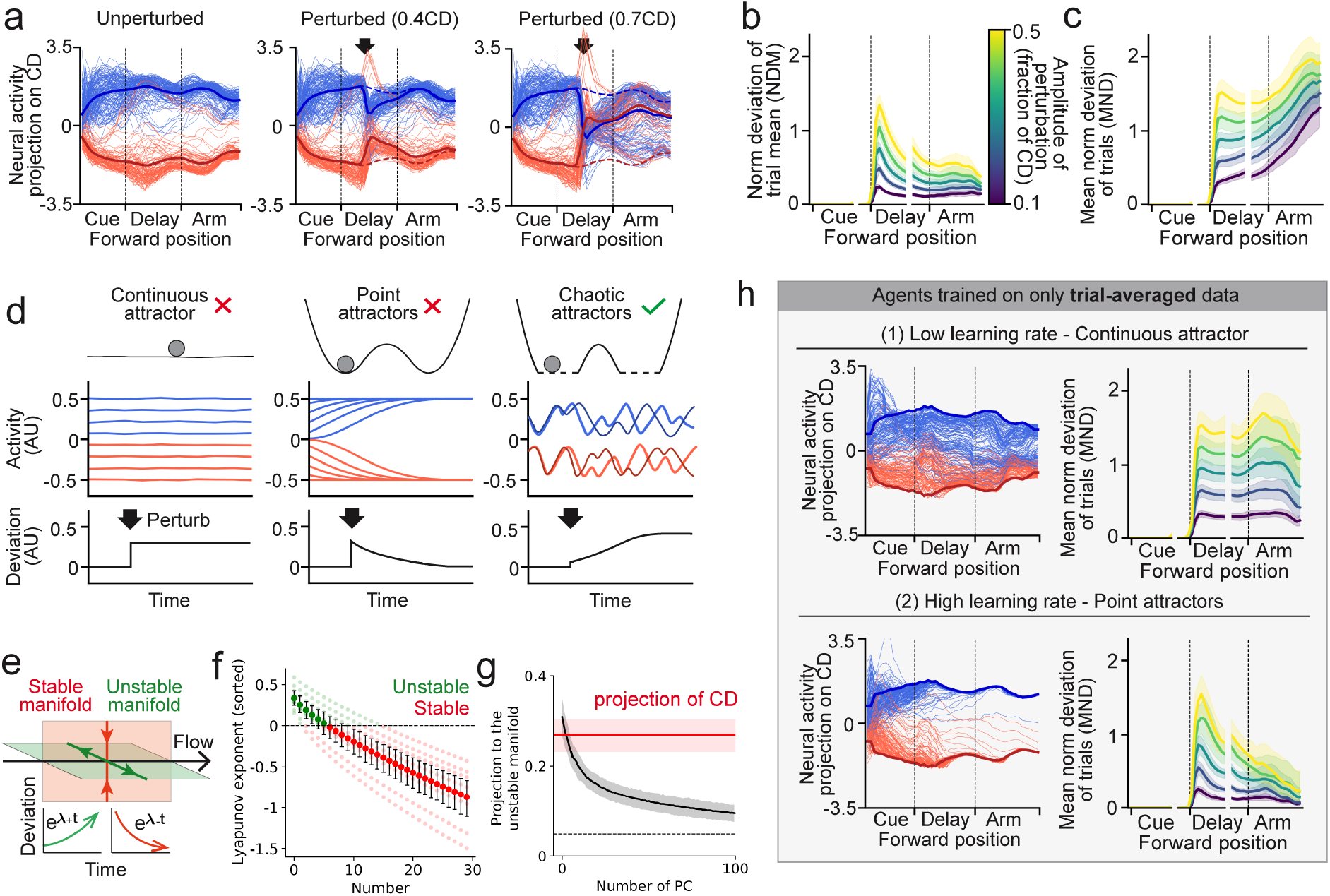
Perturbative analyses reveal low-dimensional chaotic attractors in data-derived agents. **(a)** Neural activity generated by a data-derived agent in an example session (only including correct trials when unperturbed), projected on the choice dimension (CD) of the agent. Left: Unperturbed. Middle and right: Perturbed along the CD toward the opposite trial type (amplitude 0.4 or 0.7× the difference between mean left and right CD projections). Black arrow indicates perturbation time. **(b)** Evolution of the norm deviation of the trial-mean (NDM) activity following perturbations of varying amplitudes. Deviations for left and right trial means are computed separately and then averaged. **(c)** Evolution of the mean norm deviation (MND) of individual-trial activity. In (b) and (c), only trials with correct post-perturbation choices are included, so the deviation reveals dynamics within each choice. **(d)** Schematics of different dynamical regimes. Top: Potential landscapes. In chaotic attractors dotted lines are used to indicate non-canonical landscapes. Middle: Temporal evolution of activity from different initial states. Bottom: Responses to perturbation at the black arrow. Deviation remains constant for continuous attractors, decays for point attractors, and shows bounded amplification for chaotic attractors **(e)** Decomposition of dynamics onto stable and unstable manifolds. Inset: Evolution of perturbations on the stable and unstable manifolds. **(f)** First 30 Lyapunov exponents of data-derived agents. Light dots represent agents trained on individual sessions; solid dots indicate session average. **(g)** Projection of residual-space principal components (PCs) and the CD on the unstable manifold, computed per session and averaged. Dotted line indicates chance-level projection. **(h)** Learned dynamics of agents trained on trial-averaged data (low or high learning rate, top and bottom). Left: The thick blue (red) line shows the neural trajectory simulated from the averaged initial states of left (right) trials; thin lines show neural trajectories simulated from the initial states of individual trials. Right: Responses to perturbation.

We next examined a data-derived agent’s response to perturbations on individual trials. Surprisingly, on individual trials, small perturbations caused deviations in neural activity patterns that diverged by increasing amounts over time and did not return to their original trajectories (Fig. 3c). We quantified these deviations using the mean norm deviation (MND) of individual trials. Because deviations occurred in different directions across trials, the mean network response appeared to return to the original average trajectory (Extended Data Fig. 2c). Notably, this divergence after perturbation occurred across all dimensions of neural activity, including components aligned or not aligned with the choice dimension (Extended Data Fig. 2c, d). These findings contrast with traditional attractor models, in which activity on individual trials reliably returns to the same trajectory after small perturbations (Fig. 3d). Instead, the progressive divergence of neural trajectories following a small perturbation is a hallmark of chaotic dynamics ^30,47^. Thus, our results reveal the presence of chaotic dynamics within each choice-specific attractor, motivating the concept of chaotic attractors. Intuitively, these dynamics amplify small fluctuations in neural activity over time, intrinsically generating trial-to-trial variability. However, within the broader attracting landscape, these chaotic dynamics remain confined to the basin associated with a single choice.

To further characterize the chaotic attractors, we identified normal modes of the dynamics, which are characteristic directions along which evolution of neural activity is self-sustained, i.e., independent of other directions ^48,49^. Along stable directions, infinitesimal perturbations decay exponentially, while along unstable directions they grow exponentially (Fig. 3e). The exponential growth rate along each mode is quantified by its Lyapunov exponent, and the corresponding direction is referred to as the covariant Lyapunov vector. Lyapunov exponents and covariant Lyapunov vectors can be viewed as nonlinear analogs of eigenvalues and eigenvectors, with the key distinction that the vectors vary with system state due to nonlinearity (see Methods for details).

We numerically computed the Lyapunov exponents of the data-derived agents and identified the span of the leading unstable covariant Lyapunov vectors ^48,49^ (Extended Data Fig. 2e, f). Only a small subset of directions exhibited instability, as indicated by their positive Lyapunov exponents (Fig. 3f). These unstable directions define a low-dimensional unstable manifold along which perturbations amplify over time. The directions of this manifold are state-dependent, reflecting the nonlinear nature of the underlying dynamics. Due to its nonlinear geometry, the low-dimensional unstable manifold is embedded within a much higher-dimensional neural space, consistent with amplified deviations appearing across many activity dimensions. The choice dimension projected strongly onto this unstable manifold, explaining its disproportionate share of trial-to-trial variance (Fig. 3g).

Because chaotic dynamics are closely linked to trial-to-trial variability, one might wonder whether they arose because the agents were trained to reproduce individual-trial neural activity and behavior trajectories. Indeed, training on left- and right-trial-averaged neural and behavioral trajectories did not produce chaotic dynamics. Instead, the agents learned either a continuous attractor or a pair of point attractors, depending on the learning rate (Fig. 3h). These agents trained on trial-averaged data used the same architecture and optimization as the individual-trial agents, underscoring that it is essential to consider the trial-to-trial variation to reveal the underlying dynamics.

One potential concern is whether partial, noisy observations of a stable system could be misinterpreted as chaos ^50^. Trial-to-trial variation can arise either from intrinsic chaotic dynamics or from random noise. Additionally, our empirical data sample only a subset of neurons involved in the task’s dynamics. To address this concern, we used an agent trained on trial-averaged data as the teacher. The teacher exhibited point attractor dynamics. We then generated individual trials from the teacher by adding i.i.d. Gaussian noise (Extended Data Fig. 3a). Thus, the resulting trial-to-trial variability was due entirely to injected noise, not intrinsic dynamics. We trained a student agent on trial-by-trial observations from only a fraction of the teacher’s units. Despite partial and noisy observations, the student recapitulated the teacher’s point attractor dynamics (Extended Data Fig. 3a, b). Therefore, the modeling framework can distinguish unstructured noise from chaos even with partial observations, reinforcing the evidence for chaotic dynamics in the agents trained on individual trials of mouse data.

Thus, the modeled cortical network appears to solve the navigation task using chaotic attractors, which share features with, but also differ from, stable attractors proposed previously for decision-making. Like stable attractors, the chaotic attractors segregate choices and remain robust in the presence of ongoing variability, even from chaotic dynamics. Moreover, chaotic dynamics contribute to the variability along the task-relevant dimensions and define the choice-specific dynamical reservoirs of possible navigation trajectories.

### Circuit connectivity motifs for trial-to-trial variability

We examined the connectivity of the RNN in the data-derived agents, interpreting each connection weight as a functional interaction between the corresponding real neurons, rather than as a direct anatomical synapse ^19,31^. We focused on connections between units whose activity patterns were likely important for task performance. First, we identified choice-selective units that could contribute to separating neural trajectories for the two choices (Fig. 4a, b; Extended Data Fig. 4a). Second, because trial-to-trial variability occurs in running trajectories within a trial type, we focused on choice-selective neurons that also encoded the mouse’s lateral running velocity (Fig. 4a, b; Extended Data Fig. 4b). These neurons exhibited diverse lateral velocity preferences, forming a tuning spectrum that spanned the entire lateral velocity range (Fig. 4c). Moreover, many of these neurons’ lateral velocity tuning was modulated by spatial position in the maze (Fig. 4a, b; Extended Data Fig. 4c), indicating that they represented specific actions at specific locations for a given choice. These cells provide all the components needed to mediate a choice-specific navigational trajectory. Thus, we observed neurons that encoded not only discrete decisions but also the variability in how those decisions were executed as navigational trajectories. The data-derived agents reproduced the tuning of these neurons, even in test trials withheld from training (Fig. 4d, e; Extended Data Fig. 5a–d). The ability of the data-derived agents to recapitulate single-neuron tuning further supports their biological plausibility. This, in turn, enables direct examination of the inferred connectivity among neurons with specific task-related representations.

**Fig. 4.**
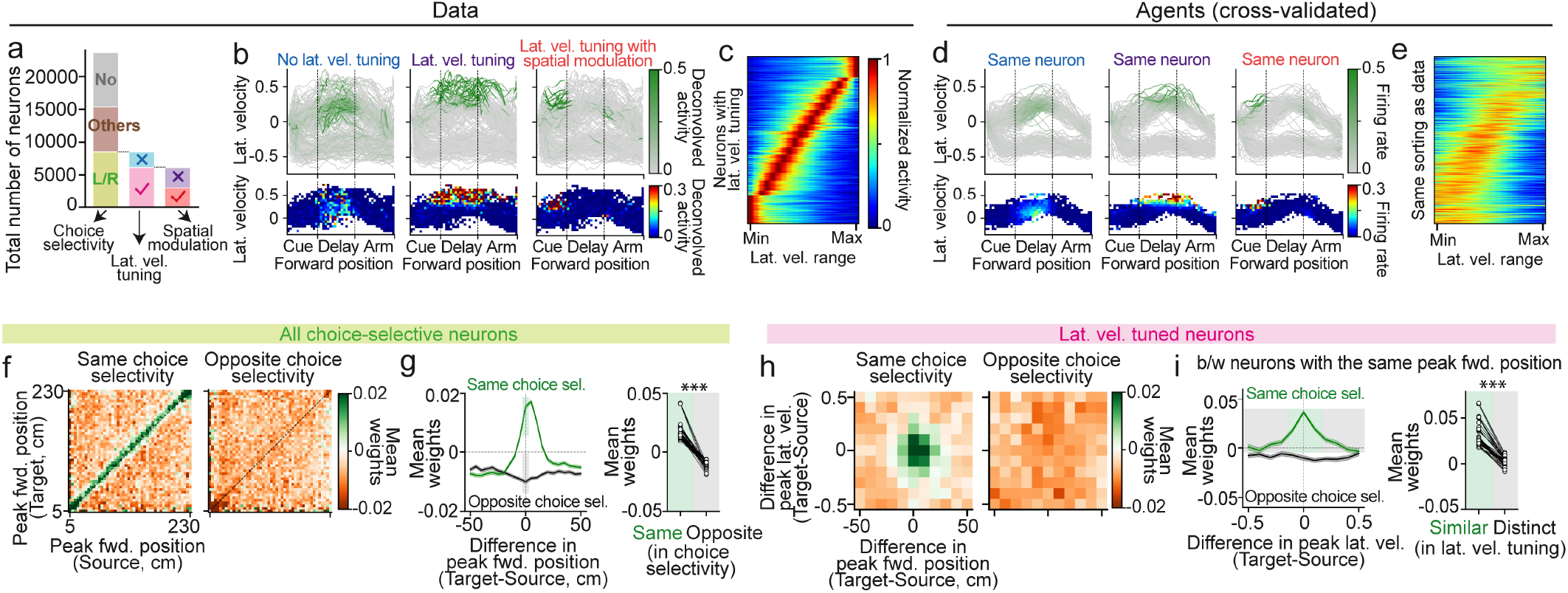
Weight analyses reveal competitive connectivity motifs on two scales. **(a)** Classification of neuron representations, pooled across areas and sessions. Neurons are first classified by choice selectivity (no selectivity, left/right selectivity, or others with multiple fields). Left or right choice-selective neurons are further classified by lateral velocity tuning, and velocity-tuned neurons are further subdivided by whether their tuning to lateral velocity is modulated by forward position in the maze. **(b)** Activity of three example left choice-selective neurons. Top: Running trajectories color-coded by instantaneous neural activity. Bottom: Bin-averaged neural activity in conjunctive bins of forward position and lateral velocity. **(c)** Neurons have diverse lateral velocity tuning that spanned the entire lateral velocity range. Activity within each neuron’s position field is plotted over its available lateral velocity range, defined by the minimum (most rightward) and maximum (most leftward) lateral velocity within the position field. Neurons are sorted by their peaks. Activity is normalized. **(d)** Corresponding RNN unit activity for the three example neurons in (b), from cross-validated test trials pooled across folds. **(e)** Lateral velocity tuning of RNN units pooled across agents, in cross-validated test trials. Units are indexed as in (c) and exhibit similar tuning as the data. **(f)** Mean connection weight between neurons with the same or opposite choice selectivity, organized by peak forward positions of source and target neurons. **(g)** Left: Mean weight as a function of the difference in peak forward position between target and source neurons. Right: Mean weight between same- or opposite-choice neurons with matched peak forward position. Dots represent individual cross-validation folds of all sessions. Shaded areas in the left panel indicate the subsets analyzed in the right panel. **(h)** Mean weight as a function of the differences in peak forward position and peak lateral velocity between target and source neurons. **(i)** Left: Mean weight between neurons with matched peak forward position, plotted against the difference in peak lateral velocity. Right: Between neurons with the same choice selectivity and matched peak forward position, mean weight is further modulated by the difference in peak lateral velocity. Shaded areas in the left panel indicate subsets analyzed in the right panel. ^***^*P* < 0.001, one-sided Wilcoxon signed-rank test.

Analysis of the RNN’s connection weights showed that choice-selective neurons formed two competing populations, with neurons of opposite choice preference inhibiting each other more strongly than neurons with the same choice preference (Fig. 4f, g). This motif is consistent with opponent inhibition, which is a hallmark of decision-making models ^8,9,51–53^. Within each choice preference, neurons tended to form stronger excitatory connections with neurons active later in the sequence than with those active earlier, supporting forward propagation of the neural trajectory (Fig. 4g) ^19,20,54^. Notably, neurons sharing the same choice and spatial selectivity could be further subdivided into competing subgroups based on their lateral velocity tuning. Within these subgroups, neurons with similar lateral velocity tuning excited each other more, whereas those with distinct tuning inhibited each other more (Fig. 4h, i). Thus, the data-derived agents’ connectivity reveals a two-scale motif: (1) opponent inhibition between neurons with opposite choice preferences, potentially supporting decision-making and choice stability in attractors; and (2) fine-grained competition within each choice population, which could generate within-choice variability.

### A two-scale competition motif leads to chaotic dynamics in a toy model

To examine how chaotic dynamics could arise in the context of the two-scale motif, we generated a “toy model” as a noise-free, 400-unit vanilla RNN with random weights (Fig. 5a). The network comprised 200 “left-choice” and 200 “right-choice” units, each receiving a constant contextual input during left or right trials, respectively (Fig. 5a). This RNN’s connectivity has a 2-by-2 block structure, where the on-diagonal blocks are connections between units of same choice and the off-diagonal blocks are between units of opposite choices. The opponent inhibition between choice populations is modeled as the off-diagonal blocks being more inhibitory than the on-diagonal blocks. On top, we added fine-grained competition within each choice population. We sorted units in each choice population according to an additional tuning parameter (which can be generic, but we will call it lateral velocity tuning for connection to the empirical results). We added stronger inhibition between units with distinct lateral velocity tuning than between units with similar tuning. In this way, the units with both the same choice preference and similar lateral velocity tuning have the most positive (or least negative) weights, and we refer to them as the within-pool weights. All the other weights are the across-pool weights. This was reflected in the connectivity matrix as diagonal ridges in the on-diagonal blocks that were less inhibitory than anywhere else (Fig. 5a). The within-pool weights and across-pool weights were sampled from two Gaussian distributions: across-pool weights were drawn from 𝒩(*σβ, σ*^2^), and within-pool weights were drawn from 𝒩(*σ*(*μ* + *β*), *σ*^2^). β controlled the mean of the across-pool weights, and μ determined the difference between the mean within-pool weights and across-pool weights (i.e., specificity of the connection). We used a positive value of μ, which made the within-pool weights more positive (or less negative) relative to the across-pool weights, generating opponent inhibition both between units of opposite choices and between units of the same choice but with distinct lateral velocity tuning (the two-scale competition motif). σ set the magnitude of the gaussian distributions, determining their shapes.

**Fig. 5.**
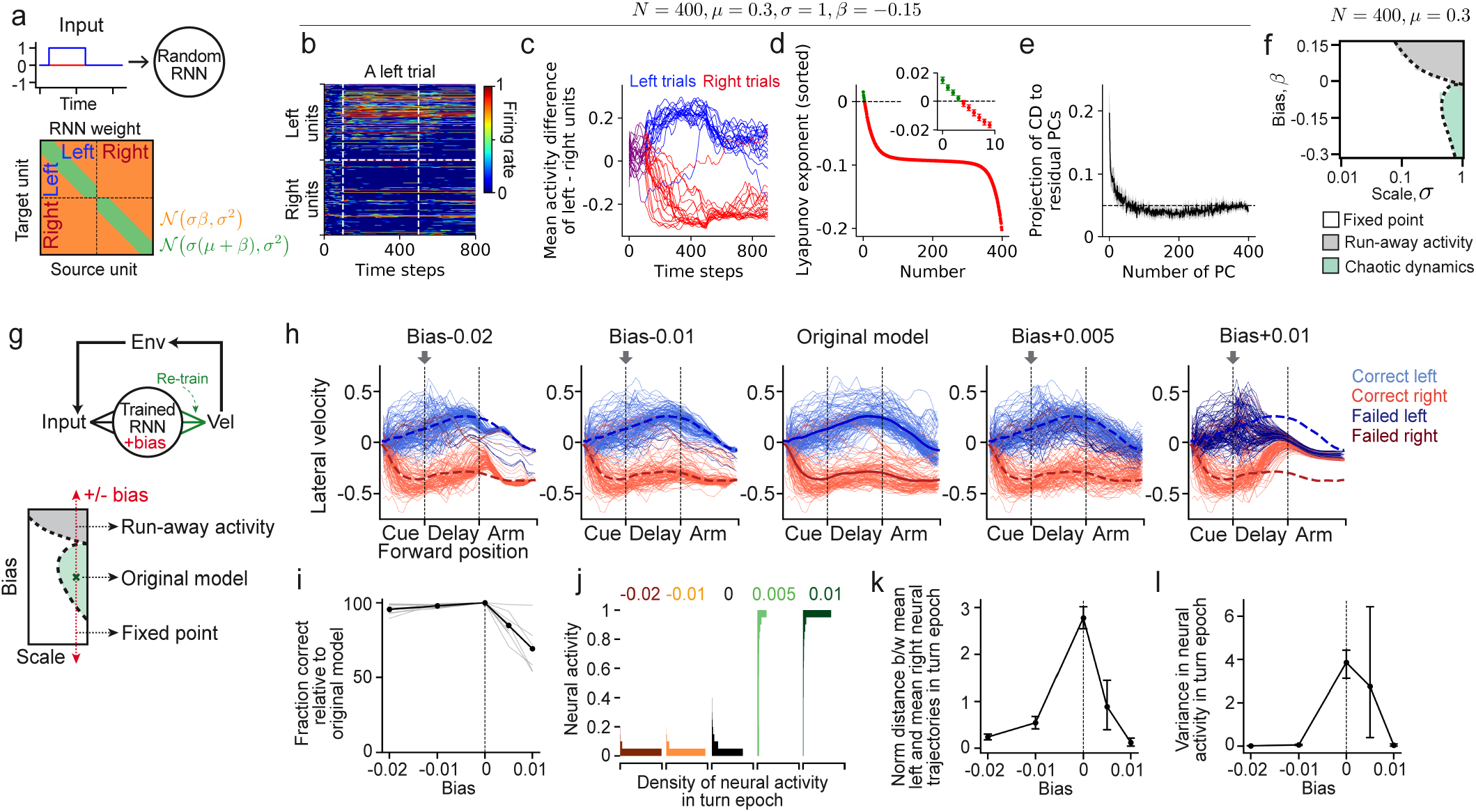
Competitive connectivity motifs and inhibitory stabilization give rise to chaotic dynamics. **(a)** Top: Schematic of the toy model. Blue and red lines represent an external input to left and right units during left trials. Bottom: Schematic of the connectivity structure of the toy model. Recurrent weights are drawn from two Gaussian distributions (green and orange partitions). Units within each choice population are ordered according to a hypothetical tuning parameter. A positive *μ* imposes competition both between choice populations (on-vs. off-diagonal blocks) and within each choice population (diagonal ridges within on-diagonal blocks). **(b-e)** Analyses of the toy model with *μ* = 0.3, σ = 1, *β* = −0.15 **(b)** Unit activity in response to the external input during a left trial. Vertical white dotted lines indicate input onset and offset. **(c)** Difference between mean activity of left and right units, in 20 left and 20 right trials simulated using identical connectivity but different activity initializations. **(d)** Lyapunov exponent spectrum of the toy model, averaged across 50 random weight realizations. **(e)** Projection of the choice dimension (CD) on principal components (PCs) of the residual space. **(f)** Phase diagram of how the dynamical regimes transition with different values of *σ* and *β*. Regimes are determined by simulations of 10 random weight realizations (Extended Data Fig. 6). **(g)** Top: Schematic of the weight perturbation procedure in dataderived agents. A bias is added to the recurrent weights at delay onset, followed by re-training only the output weights to minimize changes in velocity outputs. Bottom: Schematic of the hypothesized modulatory effect of inhibition. **(h)** Running trajectories for individual left and right trials in an example session under different weight biases. Arrow marks where the bias begins to take effect. Blue and red dotted lines indicate trial-averaged trajectories in the unmodified agent. **(i)** Choice correctness relative to the unmodified agents as a function of added bias. Thin lines represent individual sessions; the thick line shows session average. **(j-l)** Effects of inhibition level on neural activity amplitude (j), separation between left and right trial activity (k), and trial-to-trial variance of neural activity (l) in the agents during the arm epoch. Results illustrate transitions between fixed-point, chaotic and runaway regimes.

With a positive value of μ and certain choices of σ and β (μ=0.3, σ=1, β=−0.15), the simple two-scale connectivity motif strikingly led to chaotic dynamics in this toy model. Specifically, with a step input on left trials, left-choice units had higher activity than right-choice units but did not settle to a steady amplitude. Instead, they displayed complex temporal fluctuations (Fig. 5b). We summarized responses to contextual inputs as the mean activity difference between left- and right-choice units (Fig. 5c). Simulations with different random initial states produced distinct trajectories, consistent with chaotic dynamics. The resulting chaotic dynamics even had properties that matched those observed in the data-derived agents. Lyapunov exponent analysis revealed that, as in the data-derived agent, the toy model formed a low-dimensional unstable manifold, with only the first four exponents having a positive value (Fig. 5d). The leading principal components of trial-to-trial variation in the toy model were oriented toward the choice dimension, consistent with our observations in both the mouse data and the data-derived agents (Fig. 5e).

Closer examination of the toy model parameters revealed a phase transition between fixed-point, chaotic, and runaway regimes (Fig. 5f; Extended Data Fig. 6). With μ kept constant, which sets the specificity of within-versus across-pool weights, the toy model exhibited fixed-point dynamics when the weight magnitude (σ) was small. As the weight magnitude increased, dynamics transitioned from fixed points to runaway activity, characterized by saturated activity in all units in the toy model. Runaway activity could be stabilized by an inhibitory background in the connection weights (negative β). Chaotic dynamics emerged when a high weight magnitude was paired with sufficient inhibition, possibly through a nonlinear interplay between expanding and contracting dimensions. Together, these results highlight that chaotic dynamics can emerge from a combination of the two-scale competition motif and inhibition stabilization.

### Inhibition stabilization of chaotic dynamics in the data-derived agents

Consistent with previous work on inhibition-stabilized networks ^53,55–57^, the toy model suggests that inhibition plays a critical role in constraining neural instability. Particularly, there is an empirical phase transition between dynamical regimes modulated by the level of inhibition. Intermediate levels of inhibition permit chaotic dynamics, too little causes runaway activity, and high levels lead to fixed points. We tested these ideas from the toy model in our data-derived agents. We adjusted inhibition amplitude in the data-derived agents by adding a bias to all recurrent connection weights. Negative bias increased inhibition, and positive bias reduced it (Fig. 5g). To isolate transitions between dynamical regimes from mere transformations of the dynamics, we retrained the linear output weights to minimize changes in velocity outputs, reducing behavioral effects of potential rescaling, translations, or rotations in the RNN’s neural activity.

Overall, observations supported the theory of a phase transition modulated by inhibition. For the behavioral outputs of the data-derived agents, increasing overall inhibition reduced variation and eventually produced converging running trajectories, consistent with fixed point attractors. Conversely, reducing overall inhibition blurred left–right trajectory separation, eventually eliminating it, consistent with runaway activity (Fig. 5h). Reducing inhibition impaired binary choice performance far more than increasing it (Fig. 5i). For the RNN unit activity of the data-derived agents, reducing inhibition increased activity but reduced choice separability and trial-to-trial variability within a choice, consistent with runaway activity (Fig. 5j–l). Thus, the level of inhibition is a critical parameter for neural dynamics, with chaotic regimes emerging only at intermediate inhibition.

### Chaotic dynamics facilitate behavioral adaptation

Because chaotic dynamics generate variability across trials for the same decision and maintain a reservoir of possible trajectories, they may enhance behavioral adaptation, in particular to allow animals to achieve the same task goal in different ways. In navigation, the same discrete decision (turn left or right) can be achieved by many running trajectories, with changes in the environment altering which trajectories are preferable. To test if chaotic dynamics can serve this type of adaptation, we asked the data-derived agents to navigate around novel obstacles in the Y-maze to reach the goal location. Aided by the closed-loop design, the data-derived agents can be tested in environments beyond those originally used to collect the data for their training, which allows testing behavioral capacities of the data-derived neural mechanisms.

In the Y-maze, we placed two obstacles in the left arm, requiring new steering movements to reach the reward zone (Fig. 6a). The obstacles were positioned so they could not be bypassed by simply adding a bias to the velocity output. On first exposure, nearly all left-choice trials were blocked by the obstacles (Fig. 6a, middle). Then, with the RNN weights fixed, we used reinforcement learning to train only the linear output weights that translate the agent’s internal dynamics into velocity outputs ^58^ (Fig. 6c, Methods). This preserved the data-derived dynamics obtained from fitting to empirical neural activity and tested their functional capacity for learning the new task. After re-training, the agent avoided the obstacles and reached the reward location with high accuracy (Fig. 6a, bottom). To bypass the first obstacle, the agent steered more left early in the arm epoch, then steered more right later to avoid the second obstacle (Fig. 6b, top). At the same time, it steered more in trials expecting collisions and less so when far away from obstacles, avoiding overcompensation (Fig. 6b, middle and bottom).

**Fig. 6.**
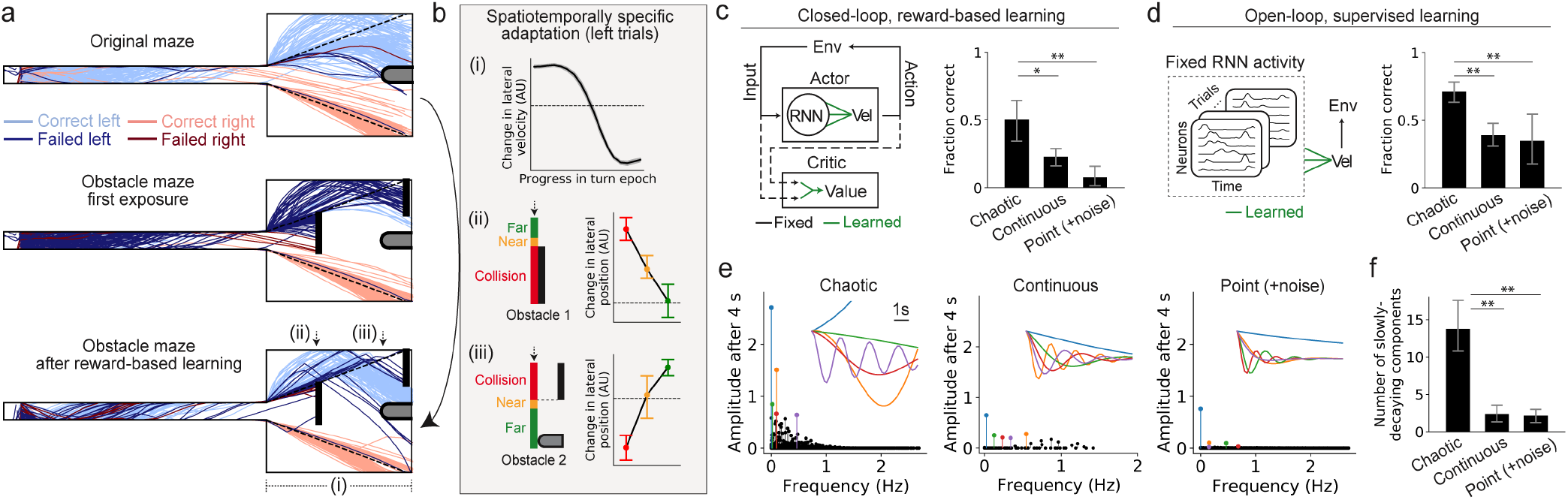
Chaotic dynamics facilitate flexible adaptation in a novel obstacle environment. **(a)** Running trajectories from an example session. Top: A data-derived agent’s running trajectories in the original maze where the neural and behavioral data used to train the agent was collected. Middle: On first exposure to the obstacle maze, most left-choice trials are blocked by the obstacles (failed trials in dark blue, obstacles shown as black vertical lines). Bottom: After reward-based re-training of only output weights, with recurrent weights held fixed, the agent avoids obstacles while maintaining high choice accuracy. **(b)** Spatiotemporally specific adaptation in the example session. Top: Change in lateral velocity (before vs. after reward-based learning) as a function of forward progress during the arm epoch, averaged across all correct left trials. Middle and bottom: Change in lateral position in front of the first or second obstacle depends on whether trajectories would have collided or were far from the obstacles. **(c)** Left: Schematic of the actor-critic algorithm used for reward-based learning. RNN weights are fixed while output weights are further trained to produce actions with higher values. Right: Performance in the obstacle maze (left trials only) compared across agents with chaotic dynamics (derived from individual-trial data) and agents with continuous or point attractor dynamics (derived from trial-averaged data). Performance is computed on cross-validated test trials and averaged across sessions. **(d)** Left: Supervised learning setup. RNN activity sequence generated by data-derived agents in the original maze is recorded and fixed while output weights are retrained to produce desired velocity outputs. Right: Performance comparison of the three agent types in the obstacle maze (left trials only) under supervised learning. Performance is computed on cross-validated test trials and averaged across sessions. **(e)** DMD eigenvalue spectra of RNN activity in agents with different dynamics (example session). The amplitudes and frequencies of components are determined by the real and imaginary part of the eigenvalues of a fitted linear operator. Insets show temporal evolution of the top five components. **(f)** Number of slowly decaying components in agents with different dynamics. Components with growth rates between −0.2 and 0.2 s^−1^ are considered as slowly decaying. Bars show session average. ^*^*P* < 0.05; ^**^*P* < 0.01, one-sided Wilcoxon signed-rank test.

To test whether this spatiotemporal adaptability is a property afforded by chaotic dynamics, we also evaluated agents pre-trained on trial-averaged data that exhibited continuous or point attractor dynamics (Fig. 3). For point-attractor agents, we added noise to match the trial-to-trial variability of the other agents for a fair comparison. After re-training with the same protocol, continuous- and point-attractor agents improved less than the agents with chaotic dynamics (Fig. 6c). Additionally, to assess the intrinsic expressivity of the neural dynamics independent of the complex learning dynamics of reinforcement learning, we took neural activity generated by the closed-loop agents in the original environment and held it fixed, and only retrained the mapping from neural activity to behavior by supervised learning. Consistently, the agents with chaotic dynamics achieved the best performance among the three types (Fig. 6d). Thus, chaotic dynamics confer greater behavioral adaptability than traditional attractors.

The benefit of chaotic dynamics to adaptability is broadly consistent with the theory of reservoir computing, in which a weakly chaotic system works as a rich reservoir of nonlinear functions that can be linearly combined to generate diverse target outputs ^59–61^. We examined this idea in the data-derived agents by applying dynamic mode decomposition (DMD) ^62^ on the RNN activity. DMD is a data-driven approach that uncovers the dominant oscillatory components of dynamical systems. The agents with chaotic dynamics had more slowly decaying oscillatory components compared to the ones with continuous or point attractors (Fig. 6e, f, Extended Data Fig. 7). In theory, by tuning the linear readout layer, these oscillatory components can be combined in various ways, with specific loading and phase offset of each component, therefore generating a large variety of outputs (see Methods for details).

## Discussion

In many sequential behaviors, animals must achieve reliable outcomes while expressing variability in the specific trajectories used to reach those outcomes. Our results identify a dynamical regime in which these two demands can coexist. We show that neural population activity can operate in a chaotic regime that remains confined within choice-specific attractor landscapes, such that trial-to-trial variability is generated intrinsically while higher-level goals remain robust. In this regime, small perturbations lead to divergent neural trajectories on individual trials, yet population activity remains constrained within a basin associated with a given choice, preserving behavioral accuracy. These findings suggest that chaos can serve a viable functional role in neural population dynamics, enabling structured variability rather than reflecting unstructured instability or noise.

This functional regime - chaotic at the level of individual trajectories yet constrained at the level of behavioral goals - can be difficult to identify using standard approaches. When neural dynamics are evaluated in open-loop settings by fitting neural activity alone or prioritizing trajectory predictability, such regimes often appear unstable, uninterpretable, or poorly predictive, and are therefore disfavored during model fitting or selection ^15,29^. In contrast, when the same dynamics are evaluated under closed-loop interaction with the environment, their importance becomes apparent: chaotic dynamics can support reliable behavior while also generating rich trial-to-trial variability. This distinction helps explain why chaotic dynamical regimes, despite being theoretically well characterized ^30,47,63^, have rarely been implicated as viable mechanisms for neural computation in behaviorally grounded models.

The contrast between open-loop and closed-loop conditions reveals closed-loop interactions as a fundamental theoretical constraint on neural dynamics. In natural behavior, neural activity, behavioral outputs, and sensory inputs are mutually coupled, such that variability in neural state necessarily reshapes future inputs and behavioral demands ^21–24^. When this coupling is ignored, the causal link between neural dynamics and behavior is broken, which can lead to systematic misclassification of which dynamical regimes are causally related to behavior. Our results show that identifying neural dynamics capable of supporting robust yet flexible behaviors over extended trajectories requires analyzing and modeling them in closed loop. Under this key constraint, dynamics that can appear unstable or uninformative in open-loop analyses instead emerge as well suited to behaviors that demand robustness at the level of goals/outcomes and flexibility at the level of trajectories.

Our findings also clarify how chaotic dynamics relate to classic dynamical frameworks that emphasize either stability or sequential structure. A long line of work has proposed that attractor dynamics can segregate neural activity into discrete choices and support robust choice formation, including in decision-making models that rely on recurrent competition and slow reverberation ^8–12^. At the same time, many models of sequential activity emphasize mechanisms that generate stable trajectories through recurrent structure and propagation ^13–15^. In the regime we identify here, these perspectives can be integrated: the global attractor landscape provides robustness at the level of the choice or goal, while within each basin, low-dimensional chaotic dynamics generate rich within-choice variability in neural trajectories. In this sense, chaotic attractors allow a single network to maintain stable high-level outcomes while supporting flexible, variable realizations of those outcomes over extended trajectories.

We also motivate a reframing of trial-to-trial variability in neural population activity. Prior studies across sensory representation, decision-making, and movement have emphasized different interpretations of variability. These include the possibility that variability aligned with coding dimensions can act as information-limiting noise correlations ^64,65^, that variability may largely lie outside task-relevant dimensions and therefore be ignorable for downstream computations ^66–70^, or that variability can reflect meaningful internal state or history dependence that supports computation over time ^1,3–5,71–73^. Here we propose a complementary perspective grounded in task demands. In sequential behaviors that admit many valid trajectories toward the same goal, variability in how the goal is realized can be an essential component of the solution rather than a nuisance. In our setting, focusing only on the trial-averaged solution risks imposing an overly rigid view of the underlying dynamics. Consistent with this, restricting training to trial-averaged neural and behavioral trajectories has classically produced point-attractor or continuous-attractor dynamics rather than chaotic dynamics, underscoring that trial-resolved variability is informative about the underlying regime and can be necessary to reveal it.

Our findings are broadly consistent with ideas from reservoir computing, which emphasize the computational capacity of rich recurrent dynamics and have highlighted “edge of chaos” regimes as potentially favorable for balancing memory and sensitivity to inputs ^29,59–61,74^. In that literature, the term “reservoir” typically refers to a large recurrent network whose high-dimensional dynamics provide a rich set of nonlinear transformations, such that only a task-specific readout needs to be trained to realize particular computations. Here we adapt the term to be more behaviorally grounded and use “dynamical reservoirs” to describe how the chaotic attractors we identify can maintain a distribution of possible navigation trajectories that can be flexibly selected, and in our simulations adapted, to meet task constraints. Importantly, the variability generated by the dynamical reservoirs here differs from unstructured noise in two ways that matter for behavior. First, chaotic dynamics are deterministic, and the nonlinear functions embedded in their dynamics can be combined to generate structured output patterns in ways that injected noise typically cannot. Second, because the chaotic dynamics remain bounded by the overall attractor landscape, they can support variability without undermining the robustness of goal-level performance that unbounded instability or noise might compromise.

The circuit structure that emerged in our data-derived agents extends commonly studied motifs in a way that naturally supports constrained variability. Consistent with canonical decision-making models and recent experimental work, our modeling revealed an opponent inhibition motif between pools of neurons with opposite choice selectivity ^8,9,51–53^. We also observed asymmetric connections that support sequence propagation, as proposed in many sequence-generation models ^13,20^. Strikingly, within a group of neurons sharing the same choice selectivity and similar sequential position, we found additional competition motifs organized by tuning to specific running trajectories, suggesting a fine-grained structure that can support within-choice variability. This type of fine-grained competition is reminiscent of that found within populations for sensory perception ^75,76^. In our toy model simulations, a two-scale competition motif coupled with inhibitory stabilization was sufficient to produce low-dimensional chaotic dynamics with bounded variability, and perturbations of inhibition in the data-derived agents produced corresponding transitions between fixed-point, chaotic, and runaway regimes. Together, these results suggest that constrained instability, and specifically chaos stabilized by inhibition, can provide a mechanistically plausible route to generating structured trial-to-trial variability while maintaining robust choice separation, consistent with broader ideas about inhibition-stabilized networks ^53,55,56,77^.

Since closed-loop coupling is a defining constraint on viable neural dynamics, we developed a novel modeling setup of data-derived agents. This approach combines data-constrained dynamical systems reconstruction ^19,31^ with environment-interacting task-performing agents ^32,41^. This setup allows modeling of behaviors that involve extended, self-paced action sequences while keeping the internal dynamics constrained by recorded neural activity and behavior on individual trials. The closed-loop design is essential because small action errors can otherwise compound into divergent environmental state distributions, and the trial-resolved fitting is essential because it preserves the structured variability that is informative about the underlying dynamics. Within this data-constrained, closed-loop modeling objective, the data-derived agents developed here can be analyzed directly as dynamical systems. They learn a generative parameterization that captures a distribution of neural trajectories rather than reproducing exact single-trial trajectories, which is expected in the presence of chaos. This framework enables controlled perturbations and dynamical analyses, such as Lyapunov spectra and unstable manifold geometry, that characterize the stability structure of the learned regime and link it to circuit-level interaction motifs. It thereby provides a tractable setting in which candidate dynamical regimes and mechanisms can be identified, probed, and used to generate experimentally testable predictions.

On the other hand, deep reinforcement learning agents, which operate in closed loop, are starting to be used in neuroscience to model behavior and representations in complex tasks ^35–40^. However, because those agents are optimized for task performance rather than constrained by trial-resolved neural population dynamics from real recordings, they address a different modeling objective: learning effective policies, rather than identifying which internal dynamical regimes are compatible with observed neural population activity during behavior.

Despite long-standing theoretical interest, the presence of chaos in the brain remains controversial, and experimentally establishing chaotic dynamics in vivo can be practically challenging ^78–82^. While our data-derived agents provide evidence that chaotic regimes are consistent with the neural and behavioral constraints in this task, directly demonstrating chaos in biological circuits will likely require perturbations and single-trial analyses that can distinguish unstable dynamics from noise. In this respect, the circuit motifs suggested by the agents and toy models, such as structured competition within choice-selective populations and inhibition-dependent regime transitions, may offer more accessible near-term targets for experimental testing than direct dynamical reconstruction alone. More broadly, our recordings span multiple cortical areas, and we modeled all recorded neurons together without area distinctions. An interesting topic is whether different areas of the cortex contribute in distinct ways to the regime we identified ^31,83,84^. In this work, we were limited in our ability to address this question because our data had imbalances in sampling across areas and limited sampling in some areas on individual sessions, likely due to inhomogeneities of imaging quality across the large cranial window (Extended Data Fig. 1a). As a result, it was challenging to address the contributions of some individual areas in some sessions and to ensure that any differences were genuine instead of due to the number of neurons from an area included in the agent. Future work with improved multi-area sampling or model structures that pool neurons across sessions could clarify whether and how different areas contribute distinctly to the dynamics.

## Acknowledgements

We thank members of the Harvey and Rajan labs for discussions and feedback on the manuscript. This work was supported by NIH grants R37NS089521 (C.D.H), R01NS143140 (C.D.H), DP1MH125776 (C.D.H.), RF1DA056403 (K.R.), U01NS136507 (K.R.), and funding from James S. McDonnell Foundation (220020466 to K.R.), Simons Foundation (Pilot Extension-00003332-02 to K.R.), McKnight Endowment Fund (K.R.), CIFAR Azrieli Global Scholar Program (K.R.), NSF (2046583 to K.R.), Harvard Medical School Neurobiology Lefler Small Grant Award (K.R.), and Harvard Medical School Dean’s Innovation Award (K.R.). This work has been made possible in part by a gift from the Chan Zuckerberg Initiative Foundation to establish the Kempner Institute for the Study of Natural and Artificial Intelligence at Harvard University.

## Author Contributions

S.Z., K.R., C.D.H conceived of the project. S.Z. developed the modeling approach, implemented all models, and performed the data analysis with guidance from K.R. and C.D.H. R.P.B. provided input on the model development and implementation. C.A. collected the experimental data with guidance from C.D.H. S.Z., K.R., and C.D.H. wrote the manuscript with feedback from all authors.

## Data Availability

Data are available upon request to the corresponding authors.

## Code Availability

All models and analyses were implemented in Python (https://www.python.org). The code to train data-derived agents with ARCTIC will be made publicly available upon publication.

---

**Extended Data Fig. 1.**
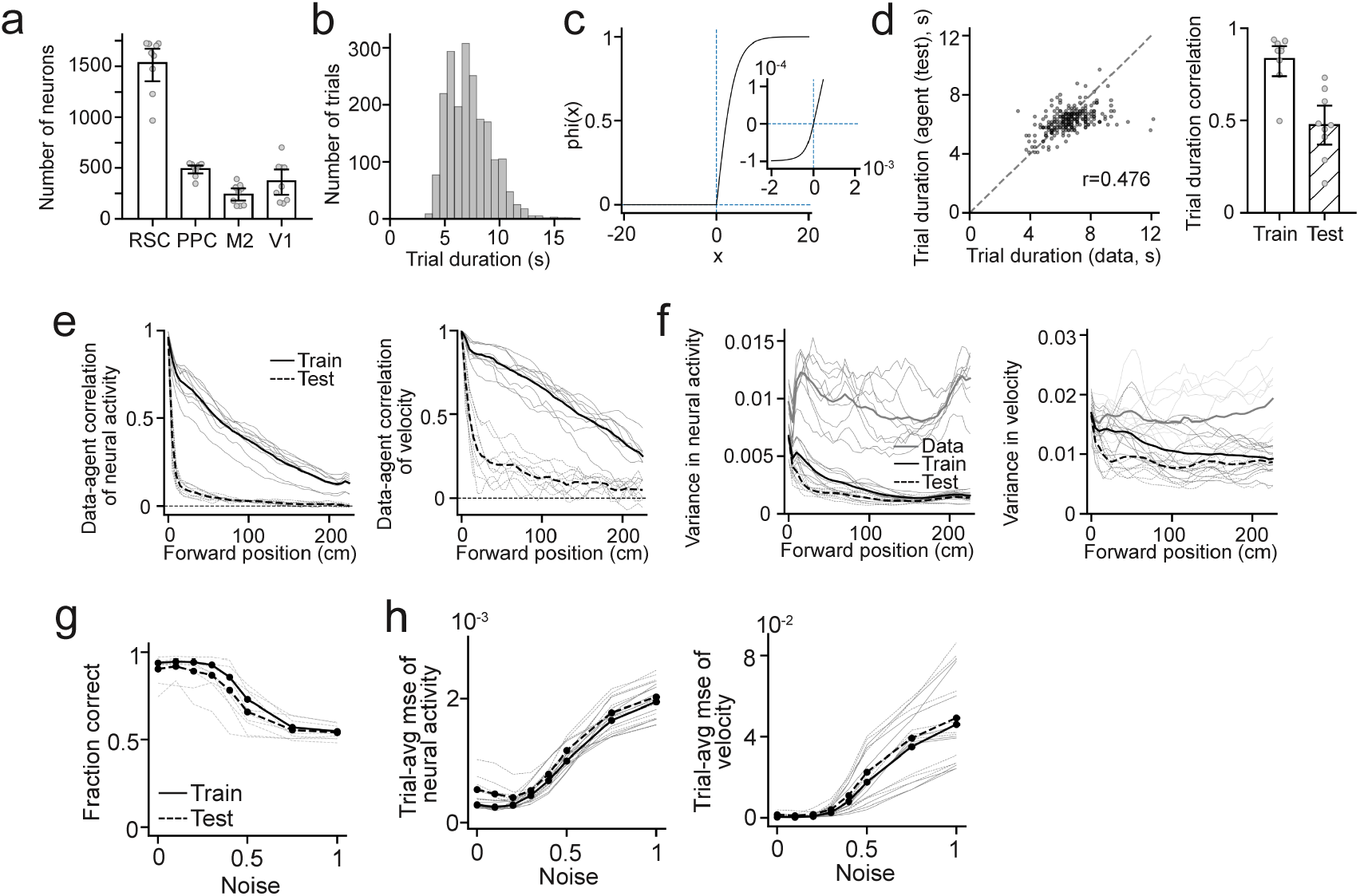
Additional data information and evaluations of the data-derived agents. (**a**) Number of neurons recorded in each of the four brain areas. Dots are individual sessions. **(b)** Distribution of the trial duration (time to traverse the maze), pooled across sessions. **(c)** Custom activation function for agent’s RNN units. Inserted panel shows the function around origin. See Methods for details. **(d)** The time taken by the data-derived agents to traverse the maze is also variable and correlates with the data. Left: Duration of cross-validated test trials from the data-derived agent plotted against the corresponding trial duration of the mouse trials in one example session. Right: Correlation of trial duration of all sessions in training and testing trials. **(e)** Residual correlation between data-derived agents and data, which reflects how the trial-to-trial variation of the agents matched that of the data. Calculated from position-binned variables. Thin lines are individual sessions, averaged across each session’s five crossvalidation folds. Thick black lines represent the average across all sessions. As discussed in the main text, it is challenging for the data-derived agents to reproduce the exact trial-by-trial trajectories, instead it learned a generalizable distribution. **(f)** Residual variance in neural activity and velocity. As in each trial the agents are initialized with the first data point of that trial, the residual variance of the agents starts at a similar level to the data and later shows some decrease. It converges to a non-zero value later in the maze. This potentially indicates that a fraction of (but not all) the variance in the data is recapturable by internal dynamics. **(g-h)** Robustness of the data-derived agents against random noise. Note that the noise level during training is 0.1. As the data-derived agents are evaluated with increasing noise, they initially stay robust in behavioral performance, trial-averaged neural activity and trial-averaged velocity for up to three times the training noise (0.3). Beyond that the agents’ performance starts to drop to chance level.

**Extended Data Fig. 2.**
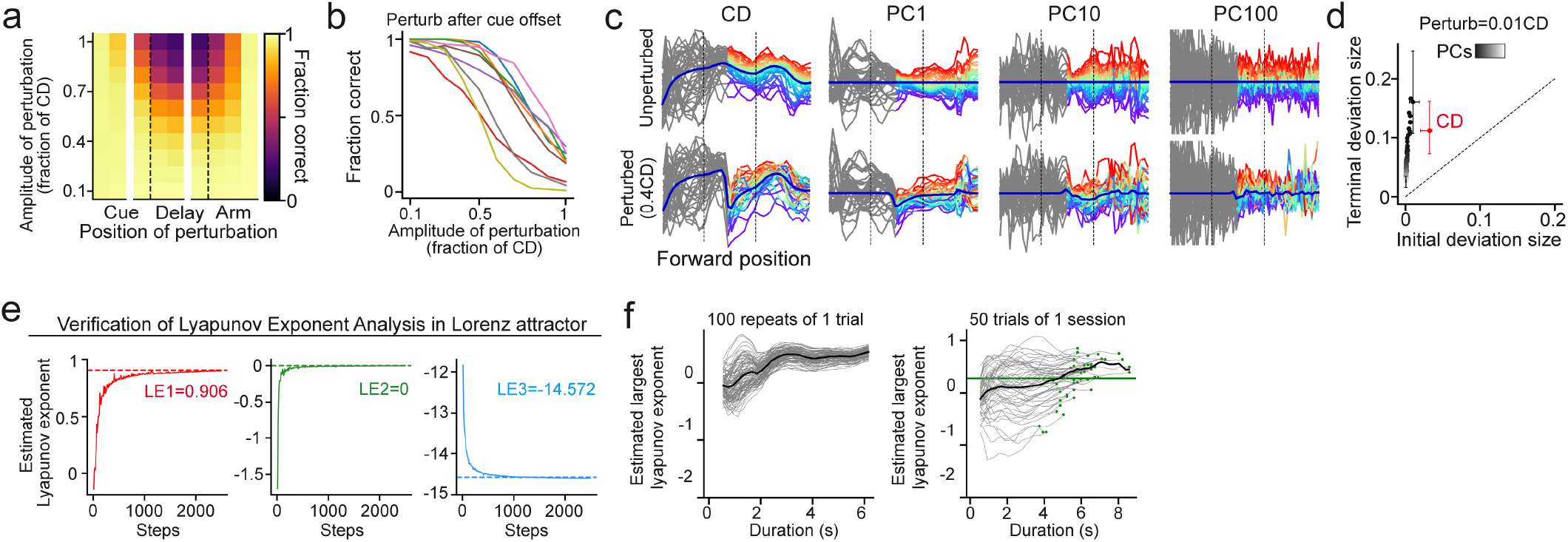
Additional results of perturbations in data-derived agents and Lyapunov exponent analysis. **(a)** Effect of perturbations of different amplitudes applied at different positions in the maze on the behavioral performance of the data-derived agents. Perturbation is more effective in the delay epoch than in the cue epoch, suggesting visual cues constrain neural dynamics into a single attractor, while during delay there are two choice-specific attractors across which a strong perturbation can drive trajectories to switch. **(b)** Perturbation effect on behavioral performance in individual sessions. **(c)** While the trial average recovers after perturbation, individual trials do not. Individual left trials in a session are colorcoded with their position-wise rank in the dimension evaluated (CD, PC1, PC10, PC100). Following perturbation, trajectories of individual trials experience shuffling (as indicated by the mixed colormap) while the trial average (thick blue curve) recovers. **(d)** Divergence occurs on all dimensions. Initial perturbation is along the CD, and evolution of the deviation on all the PC dimensions are evaluated. **(e)** Verification of the numerical method calculating Lyapunov exponents in the known system of Lorenz attractor. The dotted line marks the theoretical value of the 3 Lyapunov exponents. The curves show how the estimated values converge to the theoretical values during simulation. **(f)** For the data-derived agents, the Lyapunov exponents are estimated by running 100 repeats of 50 randomly chosen trials of each session. The estimated Lyapunov exponents show trends of convergence over the trial duration. For some trials, the estimated Lyapunov exponents might have not fully converged, making our estimation a potential lower bound. Left: Thin lines show individual repeats; the thick line shows average across all repeats for the given trial. Right: Thin lines show individual trials, thick line shows average across the 50 trials, green dots show the final estimates for each trial, and green line shows the final estimated largest Lyapunov exponent for this session, which takes the average of the estimates for individual trials.

**Extended Data Fig. 3.**
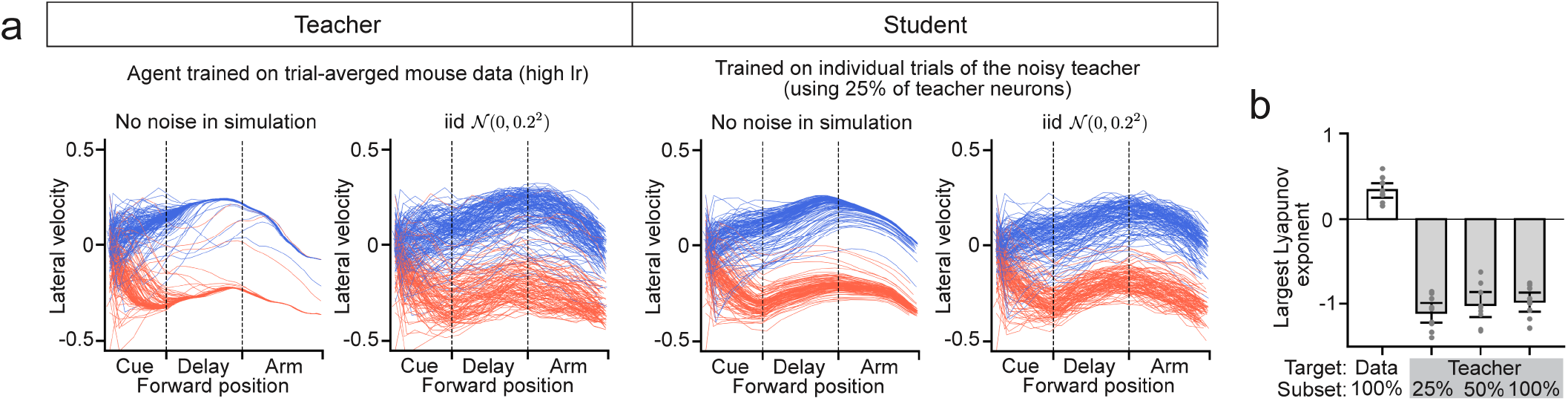
The data-derived agents distinguish unstructured noise from chaos despite partial observations. **(a)** An agent trained on trial-averaged activity and exhibited point attractor dynamics is used as the teacher. The teacher is simulated with injected unstructured noise to generate trial-to-trial variability. A student agent trained on the individual trials of the noisy teacher is able to recover that the teacher has stable dynamics, even when the student is only given a subset of teacher neurons. Figures show velocity outputs of the teacher and student agents, without or with injected random noise. **(b)** Largest Lyapunov exponent of agents fit to the individual trials of different sources (data, or a subset of neurons from the teacher). Dot are individual sessions. Negative value indicates stable dynamics.

**Extended Data Fig. 4.**
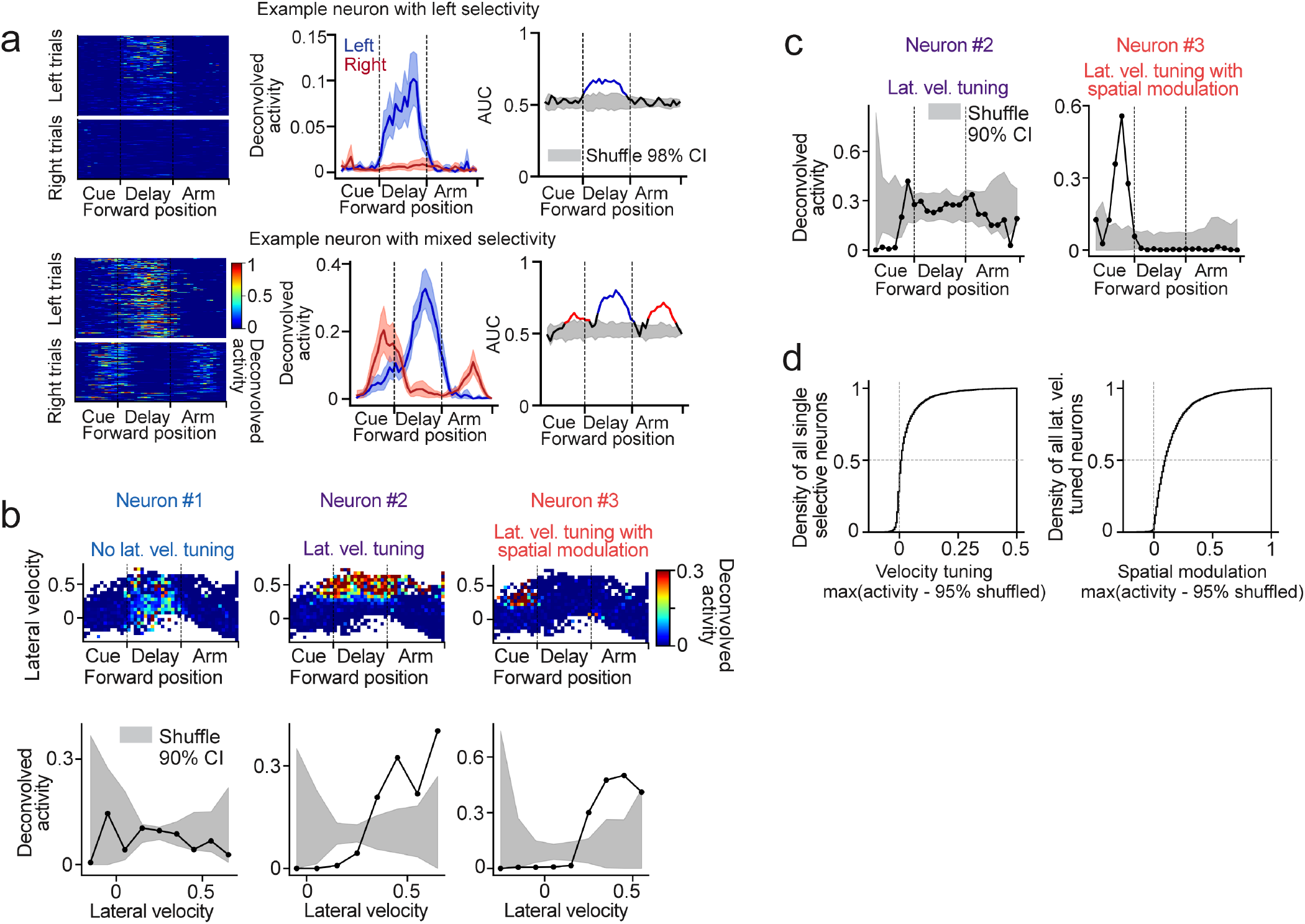
Evaluation of single neuron representations. **(a)** Two example neurons with choice selectivity. Left: Neural activity of individual left and right trials, plotted against the forward position in the maze. Middle: Activity averaged across all left or right trials. Right: Selectivity is determined by area under curve (AUC) of a support vector classifier. Shaded area shows null distribution of the AUC. Colored segments mark the selective fields of the neurons. **(b)** Lateral velocity tuning of neurons. Top: Bin-averaged neural activity in conjunctive bins of forward position and lateral velocity (Same as in Fig. 4b). Bottom: Activity around the peak forward position is plotted as a function of lateral velocity. The shuffles are generated by shuffling the velocity labels of time steps. A neuron is considered to have significant lateral velocity tuning if there is at least one lateral velocity bin where the activity exceeds 95% of the shuffled distribution. **(c)** Spatial modulation of lateral velocity tuning. Activity at time steps when the lateral velocity is within the neuron’s selective field of lateral velocity is plotted against the forward position. Neuron #3 but not #2 shows large spatial modulation. **(d)** Distribution of lateral velocity tuning and its spatial modulation. See Methods for details.

**Extended Data Fig. 5.**
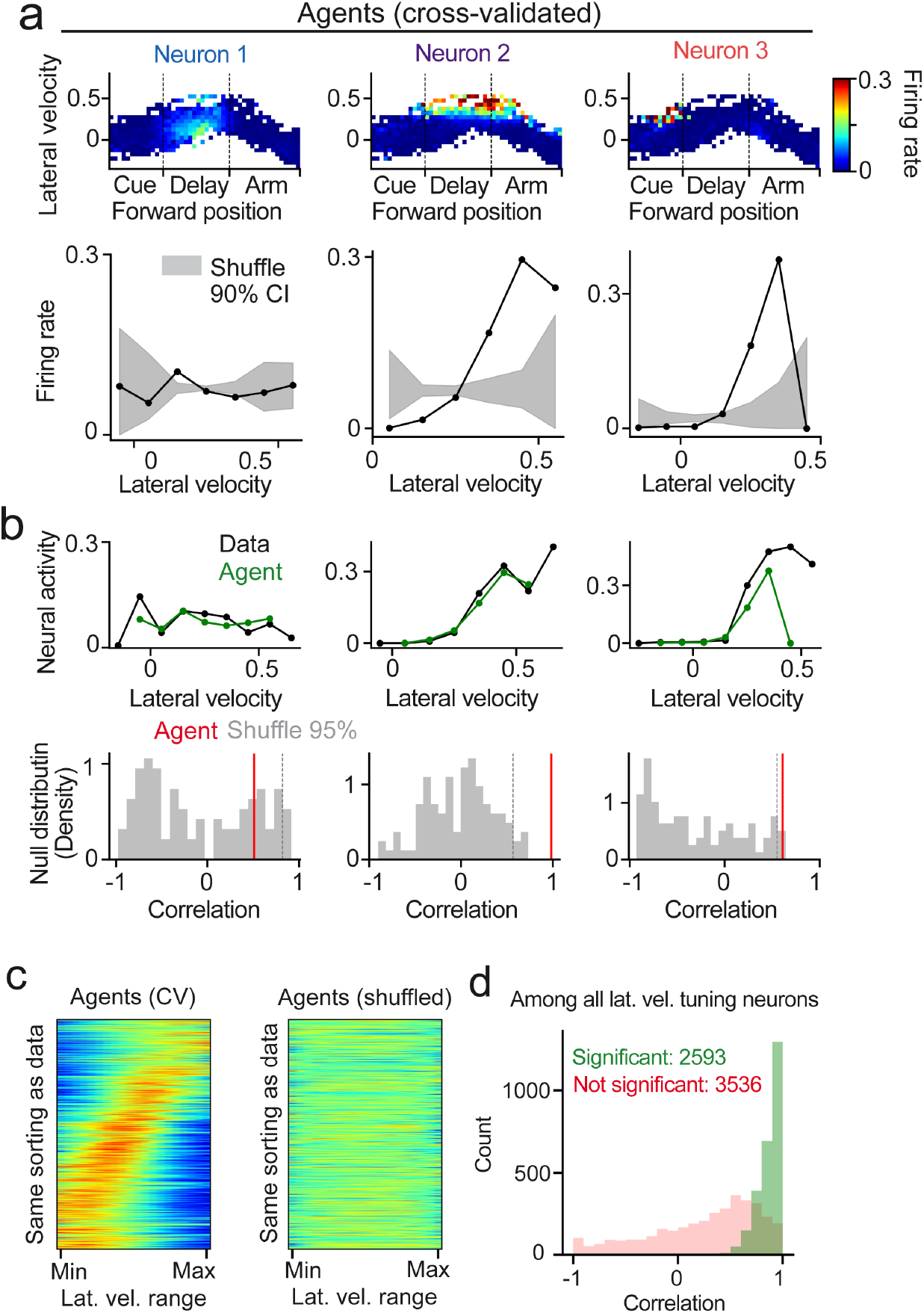
Data-derived agents recapitulate lateral velocity tuning of single neurons. **(a)** Similar to Extended Data Fig. 4b, the same analysis is repeated for the corresponding units in a data-derived agent to quantify their lateral velocity tuning. In the bottom row, the black curve is the tuning curve of the RNN unit in the data-derived agent, and the shade is the 90% CI of the shuffled tuning curves. **(b)** To evaluate how well the data-derived agents recapitulate lateral velocity tuning, correlation between the tuning curves of the data and the agents is computed. The distribution of shuffled correlation is calculated based on shuffled tuning curves. The fitting is considered significant if the correlation is greater than 95% of the null distribution. **(c)** Lateral velocity tuning generated by the data-derived agents forms a spectrum over the lateral velocity range. This feature disappears in the shuffled tuning curve, suggesting it is not due to trivial reasons (e.g., correlation between lateral velocity and forward position). Neurons are sorted with the same indexing as in the data (Fig. 3c) **(d)** Distribution of the data-agent correlation of lateral velocity tuning curves.

**Extended Data Fig. 6.**
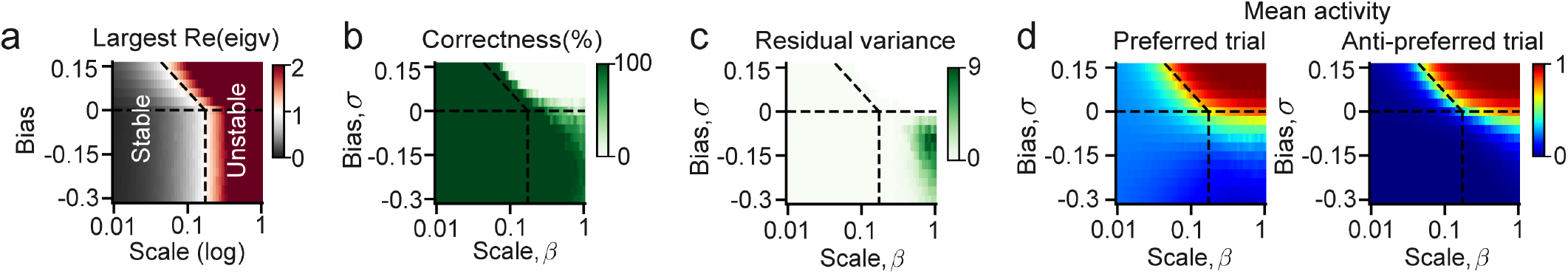
Evidence for the empirical phase transition in Fig 4f. **(a)** Largest eigenvalue of the connection matrix. Values greater than one suggest instability. **(b)** Left/right classification correctness, evaluated by whether the left or right neurons have greater activity. **(c)** Residual variance in neural activity. **(d)** Mean activity of all units in their preferred or anti-preferred trials. Note that (a-d) show results averaged across 10 random weights realization. (b-d) show average across 100 left and 100 right trials with random activity initializations.

**Extended Data Fig. 7.**
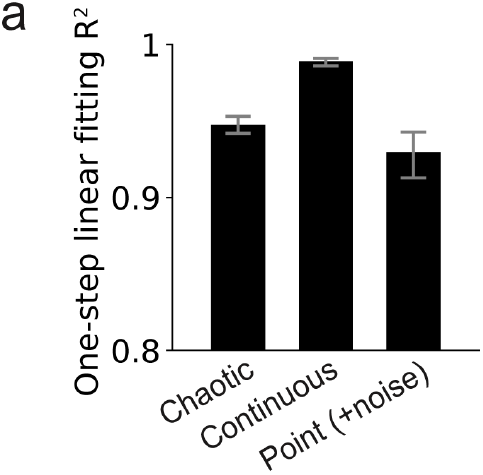
Additional information on dynamic mode decomposition (DMD) **(a)** One-step fitting performance of the linear operator in DMD, in R^2^, for the RNN activity sequence in the data-derived agents with different dynamics. Averaged across all sessions.

## Methods

### Multi-region calcium imaging in mice

All procedures were approved by the Harvard Medical School Institutional Animal Care and Use Committee and were performed in compliance with the Guide for the Care and Use of Laboratory Animals. Detailed experimental procedures was described in reference ^43^, where the data was previously reported. In brief, two female C57BL/6J-Tg (Thy1-GCaMP6s) GP4.3Dkim/J mice (The Jackson Laboratory, stock 024275) were trained to navigate in a virtual reality maze operated in VirRMEn (Virtual Reality Mouse Engine) ^85, 86^. Mice were head-fixed atop an 8-inch Styrofoam ball serving as a spherical treadmill. Movements of the treadmill were measured and converted into pitch, roll, and yaw velocities, which controlled the mouse’s forward and lateral translocation in the virtual environment. The virtual environment was projected by a micro projector onto a 15-inch diameter half-cylindrical screen. Mice were trained in a Y-maze. The total length of the Y-maze was 250 cm, with a 150-cm-long, 20-cm-wide Y-stem corridor, followed by a 100-cm-long, 80-cm-wide Y-arm funnel. The virtual location of mice could not get within 5 cm of the walls, and the virtual view angle of mice was fixed. In all trials, one of two visual cues was shown on the wall, with a consistent association between each cue and its corresponding rewarded arm in the Y-maze. Horizontal gratings signaled a left reward, whereas vertical gratings signaled a right reward. The visual cue was presented only during the first part of the maze, followed by a delay period where a neutral visual pattern not predictive of the choice was shown on the wall. For sessions used in this paper, the delay onset ranges from 70 to 100 cm in the Y-stem.

Mice were fitted with a chronic cranial window implant, exposing the dorsal surface of both cortical hemispheres ^87^ or only the left hemisphere. Data were acquired with a large field of view two-photon microscope assembled as reported previously ^44^, which allows random access imaging across a 5-mm-diameter field with 1 mm depth. Four regions (primary visual cortex, posterior parietal cortex, retrosplenial cortex, and secondary motor cortex) in the left cortical hemisphere were imaged and identified by retino-topic mapping. For each region, imaging was performed in layer 2/3 across two 600 *µ*m × 600 *µ*m planes (with a resolution of 512 × 512 pixels) separated by 50 *µ*m in depth, at 5.36 Hz per plane. We performed motion correction on the original images ^75^ and used Suite2P to extract the regions of interest ^88^. A custom convolutional neural network implemented in MATLAB was used to classify somatic sources ^75^. The mean fluorescence for each region of interest was calculated and transformed into a normalized change in fluorescence (Δ*F/F*) and then deconvolved by the constrained OASIS AR1 method ^88^. We used the deconvolved activity for all subsequent analyses.

### Data-derived agent

#### Model equation

The agent involves a recurrent neural network (RNN) interacting with a task environment in closed loop. The RNN consists of *N* unit, where *N* is chosen to match the number of neurons recorded from the mouse cortex in a given session. The state of each unit is represented by a variable *x*_*i*_, and the dynamics of the RNN is defined as

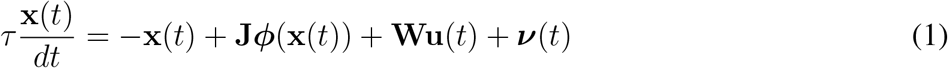

where **x** ∈ ℝ^*N*^ denotes the states of all RNN units, **u** ∈ ℝ^*E*^ denotes the environment observations, **ν** ∈ ℝ^*N*^ is a per-unit random noise drawn from an iid Gaussian distribution 𝒩 (0, *σ*^2^), **J** ∈ ℝ^*N ×N*^ is the recurrent connection between RNN units, and **W** ∈ ℝ^*N ×E*^ is the weight of a linear input layer from the environment to the RNN. *ϕ*(·) is a nonlinear activation function and *r* = *ϕ*(*x*) gives the firing rate of units. For biological plausibility ^90^, we designed *ϕ*(·) to be a modified tanh function controlled by two hyper-parameters, *r*_0_ = 0.0001, *r*1 = 4 (Supplementary Figure 1c). In this way, the firing rate is modulated to be between −*r*_0_ and 1, while the resting firing rate (when a unit receives no input from the other units nor the environment) is 0. The negative regime was further modulated by a function *g*(*x*) = *x/*(1 − 500*x*) for better numerical stability. *g*(*x*) extends the regime of negative *x* that does not suffer from numerical saturation. In practice, this helped to make *ϕ*(·) invertible on its full domain that is used in data fitting.

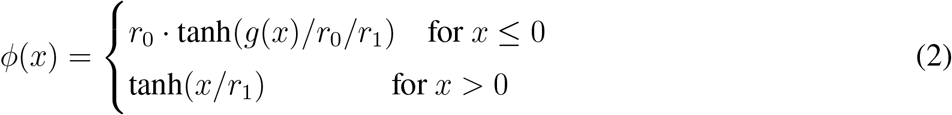

#### Environment

The environment observations are composed of three components: locomotion velocities **v**, positions in the maze **p**, and an abstracted visual cue **c**, i.e., **u**(*t*) =[ **v**(*t*) **p**(*t*) **c**(*t*)]^⊤^. **Velocity: v**(*t*) = [*v*_*f*_ (*t*) *v*_*l*_(*t*) *v*_*y*_(*t*)] are the forward(pitch), lateral(roll) and yaw locomotion velocity at time *t*, generated from the RNN via a linear output layer: **v**(*t*) = **W**^out^***ϕ***(**x**(*t*)). **Position: p**(*t*) = [**r**(*p*_*f*_ (*t*)) *p*_*l*_(*t*)] are the basis-expanded forward position and lateral position in the maze. The position is updated in the environment by integrating the locomotion velocity generated by the RNN, 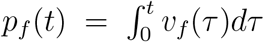 and 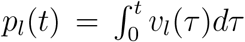. In this way, the RNN agent interacts with the environment in a closed loop. For biological plausibility, we used a set of basis functions *p*_*f*_ (*t*) → **r**(*p*_*f*_ (*t*)) : ℝ → ℝ^*M*^ to mimic how *M* idealized place cells would represent the forward position. The basis functions are designed to be cosine bumps with uniform spacing and a width-to-spacing ratio of 2. The cosine bumps are 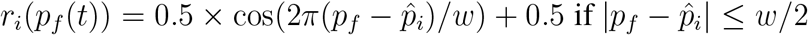 else 0, where 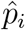 is the preferred position of the i-th “place cell” and *w* is the width of the field. We chose *M* = 5. **Visual cue:** For simplicity we abstracted the visual cue observed from the environment as a binarized variable **c** ∈ {0, 1}^2^, where **c** = [1 0] if in the cue epoch and the cue is vertical bar, **c** = [0 1] if in the cue epoch and the cue is horizontal bar, and **c** = [0 0] if not in the cue epoch. In practice, all environment observations are rescaled so their amplitudes are approximately 1.

### ARCTIC (Activity ReConstruction in Closed loop) via online imitation learning

#### Training procedure and loss

We trained one agent for each cross validation (CV) of each mouse session. The agent features an RNN with the same number of neurons as in the calcium imaging recording, and the RNN units are one-to-one matched with the real neurons. The input (**W**), recurrent (**J**) and output weights (**W**^out^) of the agent are all initialized to **0** and are then trained to reproduce single-unit activity and velocity outputs on a trial-by-trial basis. More precisely, for any trial *k* in this session, suppose it takes 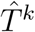 time steps to finish (reaching the end the the maze), we use 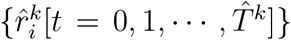 to represent the ground truth de-convolved activity of neuron *i* (*i* = 1, · · ·, *N*) and 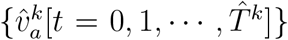 (*a* = forward, lateral and yaw) for the ground truth velocity in this trial. Then the purpose of training is if we initialize the agent’s unit activity *r*_*i*_[*t* = 0] to be the initial ground truth neural activity, 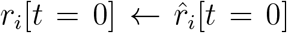, and initialize the environment such that **v**[*t* = 0], **p**[*t* = 0] and **c**[*t* = 0] all become the initial environment seen by the mouse, then starting from *t* = 0 the agent would autonomously generate an entire sequence of 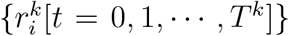 and 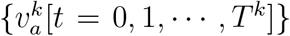 such that both the difference of the neural trajectories 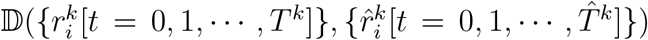 and the difference of the velocity trajectories 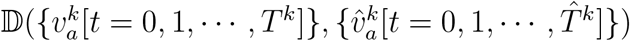 are minimized for all *i* and *a*. Because the trial duration 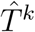 varies a lot among different trials (Supplementary Figure 1b), it is challenging for the agent to reproduce the trial duration precisely for all the trials, so that *T* ^*k*^ and 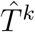 are usually not exactly the same (but can be correlated, see Supplementary Figure 1d). Therefore, it would make less sense if the loss is defined per time step as 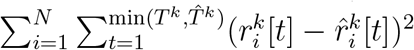. Instead, we took the idea from dynamic time warping (DTW) and aligned the trajectories to the forward progress in the maze, and we compared the aligned trajectories for the fitting error.

We did 5-fold CV. For each CV fold the training set contains 80% of all the correct trials in a session. We trained the agent for 30 epochs, where for each epoch the sequence of the training trials presented is shuffled. For each trial, the agent and environment are initialized to the first data-point of this trial. As the agent simulates forward, the agent weights are updated for each time step *t* (online learning ^42^). At each agent simulation time *t*, an instantaneous fitting error is evaluated and the weights are update accordingly. To evaluate the fitting error at agent time *t*, we need to find a target data time 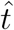 with which the agent trajectory and the data trajectory are aligned according to the forward progress in the maze. We define the target data time 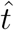 as 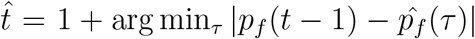. The fitting error is then computed as 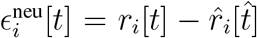 (neural error) and 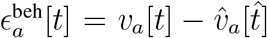 (behavioral error). Note that the weight update at agent simulation time *t* would be immediately in effect for the simulation from time *t* onward, so that it timely affected the agent’s internal activity, behavioral outputs, and resulting environmental inputs at time *t* + 1.

#### Optimization method

To optimize both the instantaneous neural error 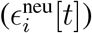 and the behavioral error 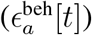 at the same time, we use the recursive least square (RLS) method as in ^19, 29^, which is a gradient-free online learning algorithm. Consider all weights that go into the RNN units as **W**^neu^ = [**J W]** ∈ ℝ^*N ×*(*N* +*E*)^, where *N* is the number of RNN units and *E* is the number of environment observations, then the update of **W**^*neu*^ follows:

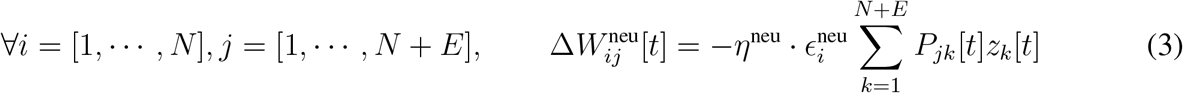

where *η*^neu^ is the learning rate. **z**^⊤^[*t*] = ***ϕ***(**x**[*t*])^⊤^ **u**[*t*]^⊤^ is the concatenation of the firing rate of all RNN units and the environment observations, which can be regarded as the “presynaptic” activity of the weights **W**^neu^. **P**[*t*] ∈ ℝ ^(*N* +*E*)*×*(*N* +*E*)^ is the inverse of the online correlation matrix of **z**[*t*], that is, **P**[*t*] = **C**[*t*]^*−*1^ and **C**[*t*] = **C**[*t* − 1] + **z**[*t*]**z**[*t*]^⊤^. In practice, instead of performing the computationally heavy inversion of the correlation matrix **C** for every single step, we use the Matrix Inverse Lemma:

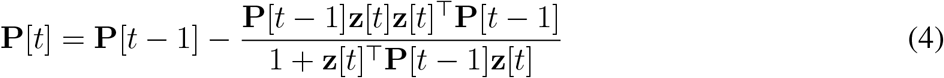

This algorithm requires **P** to be initialized properly, and we follow the convention to set **P**[0] as a diagonal matrix **I***/α*. It turns out that *α* imposes an L2 regularization on **W**^*new*^ with strength *α*. In practice, we would like to acquire a solution in which the contribution from RNN units and that from the environment are of a similar scale. We did this by applying extra regularization to the environment observations through **P**[0] = **AI**, where **A** = Diag([*α*_1_, · · ·, *α*_*N*+*E*_]), *α*_1_ = · · · = *α*_*N*_ = 1, *α*_*N*+1_ = · · · = *α*_*N*+*E*_ = 500.

Another modification we did to the conventional method is that we intentionally removed self-projections of the RNN units. Autapses (self-synapses) are relatively rare *in vivo*, yet data-derived RNN models tend to learn an unproportionally strong self-inhibition. For better biological plausibility, we fixed all self-projections to zero during training. We found that following the conventional update rule (3) and then resetting self-projections to 0 after each weight update significantly compromised the performance. Instead, we calculate a separate matrix **P**_*i*_ for each unit *i* that only tracks the non-self inputs. An efficient algorithm to achieve this is presented in the Supplementary Text.

Finally, the velocity output weights, **W**^out^ ∈ ℝ^*V ×N*^ where *V* is the number of velocity output channels, are updated using a similar rule

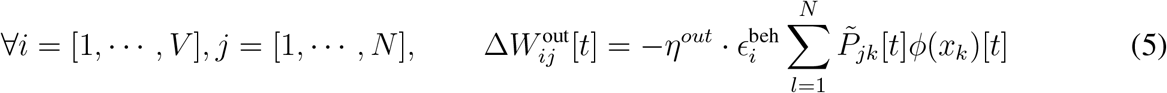

where 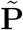 tracks the inverse correlation matrix of ***ϕ***(**x**)[*t*].

### Statistics

All error bars in the figures represented 95% confidence interval (CI) of the mean unless otherwise mentioned. The CI was calculated by bootstrapping. For a set of measurements with size *N*, 1000 random samples of size *N* were drawn from the set of measurements with replacement. The mean of each random sample was calculated, forming a distribution of the mean. The lower and upper bounds of the 95% CI of the mean were then calculated as the 2.5% and 97.5% percentile of this distribution.

### Neural representations

For population representation, choice dimension was computed per forward position bins of 5 cm. In each position bin, mean population activity of all correct left trials and all correct right trials were calculated and their difference (left-right) was defined as the choice dimension associated with that position.

The representations of individual neurons were considered in figure 4 and supplementary figure 4, which were later used to understand the connectivity motifs. The methods of computing single neuron representations are detailed below:

#### Choice selectivity

Neural activity of each trial was binned into 5 cm bins according to the forward position in the maze. Choice-selectivity only considered correct trials. Significance of choice selectivity was calculated based on AUC (Area Under the Curve). For each session, as there were usually different numbers of left and right trials, the trial-type with more trials was down-sampled randomly such that the left and right trials used for classification were balanced. A Support Vector Classifier was trained per position bin to predict the choice based on single neuron’s activity in that position bin. An ROC (Receiver-operating characterization) curve was constructed and the area under this curve was used to quantify the classifier’s performance. To compute the null distribution, the choice labels were shuffled among trials. Both the original and shuffled AUC were calculated for 100 repeats, with random sub-sampling of trials. A neuron was considered to have significant prediction of choice in a given position bin if the averaged original AUC was greater than 99% of the shuffled distribution. A neuron was then considered to have choice selectivity in a position field if (1) the neuron had significant prediction of choice in a consecutive field of at least 5 bins, and (2) the neuron had increased activity in that field, where period of increased activity was defined as the positions where the trial-averaged activity of a given choice is greater than 0.01. Neurons with left or right selectivity were defined as having only one position field selective for either the left or right choice. Neurons with multiple choice-selective position fields, whether those were for the same choice or difference choices, were classified as “others”. We focused on neurons with left or right choice selectivity in the following analyses.

#### Lateral velocity selectivity

For simplicity, lateral velocity selectivity was only considered for neurons with left or right choice selectivity. For a left (right) selective neuron, we considered all time points in left (right) trials within a 25 cm window centered at its peak forward position. The neuron’s activity was bin-averaged in conjunctive bins of forward position and lateral velocity, with the bin size of forward position being 5 cm and that of lateral velocity being 0.1. The neuron’s lateral velocity tuning curve was obtained by averaging out the forward position axis. To compute the shuffle distribution, the lateral velocity was shuffled across all time points we considered, and the tuning curve was computed according to the shuffled lateral velocity. This was repeated 1000 times to get the distribution of the shuffled tuning curve. A neuron was considered to have significant lateral velocity tuning if there was at least one lateral velocity bin where the original activity was greater than the 95% of the shuffled curves.

#### Spatial modulation of lateral velocity tuning

Spatial modulation was considered for neurons that have significant lateral velocity selectivity. We took a similar approach as above. For each neuron with lateral velocity selectivity, we considered all time points when the lateral velocity was in the neuron’s selective window, defined as lateral velocity bins where the activity was greater than the 95% of the shuffle. Activity was then binned into conjunctive bins of forward position and lateral velocity, where we used 10 cm forward position bins here. We shuffled the forward position across time points to get the null distribution. The modulatory effect of forward position was calculated by averaging-out the lateral velocity axis. A lateral velocity tuned neuron was considered to be modulated by forward position if there was at least one forward position bin where the original activity was greater than the 95% of the shuffled curves by at least 0.1.

### Agent evaluation

We claimed that the data-derived agents learned a distribution of neural trajectories from the training set and generalized such distribution to the test set. In figure 2m-q we evaluated such distribution on the population level through position-wise residual variance structures. In figure 2l, figure 4 and supplementary figure 5, we evaluated how well the agents recapitulated single neuron representations, which can be interpreted as a conditional distribution of neural activity given specific behavioral variables. In both cases, we treated each mouse session with 5-fold CV, and assembled the test trials from every fold to recover the original number of trials per session for evaluation.

For analyses of the residual variance structure, we started by binning neural activity in each trial according to the forward position in the maze into 5 cm bins. We then computed the mean population activity of all correct left or right trials. The mean activity at each position bin was subtracted from each individual trial, isolating the trial-to-trial neural activity residuals (we only considered correct trials). The total residual variance took the mean square of the residuals, summed over all neurons and averaged across position bins. We projected the residuals on choice dimension by taking the inner product, and the residual variance on choice dimension took the mean square of the projected residuals, averaged across position bins. Projected variance at chance level was calculated by dividing the total residual variance by the rank of the residuals, which was the number of correct trials as it was always smaller than the number of neurons in our data. Residual principal components were then calculated per position bins. The projection of choice dimension on each residual principal component took their inner product, and we then took the quadratic mean across position bins. The chance level projection took the square root of 1 over the rank of the residuals (number of correct trials). Evaluation of the data-derived agent took a similar procedure, while we only considered the trials where the agent made a correct choice.

For evaluation of single neuron representations, we focused on the neurons’ tuning of choice and lateral velocity. Choice selective neurons had spatially confined activity bumps, which could be characterized by a neuron’s trial-averaged activity. We sorted the neurons preferring left or right choice according to their peak forward positions, resulting in the characteristic choice-selective sequences. The same sequences were observed in the agents (Fig. 2l). The tuning to lateral velocity closely related to the trial-to-trial variation. We first visualized the lateral velocity tuning spectrum of the population. Each neuron’s lateral velocity tuning curve had the range of the possible lateral velocity visited by the mouse or the agent in the 25 cm window centered at this neuron’s peak forward position. For visualization purpose, the tuning curves were interpolated to all have 20 bins, and were normalized for each neuron. The neurons were sorted according to which one of the 20 bins had the peak activity in their tuning curves. We also compared the lateral velocity tuning curve of single neurons in the data and in the agent, by taking their Pearson correlation coefficient. A null distribution was generated by correlating the data tuning curve with the shuffled tuning curve from the agent. A neuron’s lateral velocity tuning was considered to be significantly recapitulated by the agent if the true Pearson correlation coefficient was greater than 95% of the null distribution.

### In-model perturbation

To perturb the neural activity in the data-derived agents, we added an instantaneous vector deviation to the RNN activity in individual trials at a certain forward position, chosen from [5, 30, 55, 80, 105, 130, 155, 180, 205] (in cm). The perturbation vector was chosen to be the position-specific choice dimension. The direction of perturbation was towards the opposite trial type, such that on a left (right) trial, the perturbation moved the RNN activity closer to that of the right (left) trials. The amplitude of perturbation was defined as fractions of the norm of the difference between mean left and mean right trial activity, chosen from [0.01, 0.1, 0.2, 0.3, 0.4, 0.5, 0.6, 0.7, 0.8, 0.9, 1.0], such that a perturbation of size 0.5 means that we moved the RNN activity to roughly the separation plane of the mean left and mean right trial activity. The effect of perturbation was evaluated using two metrics quantifying the deviation of perturbed neural trajectory, aligned to forward position. The first metric is norm deviation of trial mean,

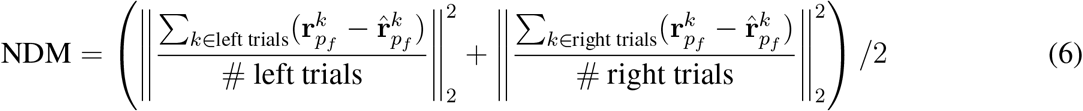

with 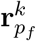 denotes the original agent activity in trial *k* at forward position *p*_*f*_ and 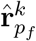 denotes the perturbed agent activity. The second metric is mean norm deviation of individual trials,

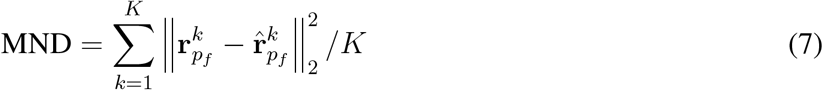

### Lyapunov exponents (LEs) and covariant Lyapunov vectors (CLVs)

In a dynamical system, the exponential growth rates of infinitesimal perturbations in different directions in the state space are described by the Lyapunov exponent (LE) spectrum, *λ*_1_, …, *λ*_*n*_ (sorted from highest to lowest) ^48, 49^. Positive and negative LEs are associated with the unstable and stable manifolds, whose directions are indicated by the covariant Lyapunov vectors (CLVs), ***γ***_1_, …, ***γ***_*n*_.

It is worth pointing out that LEs and CLVs can be viewed as a nonlinear equivalent of the log-eigenvalues (*σ*_*i*_) and eigenvectors (**v**_*i*_) of a linear operator, with the difference that CLVs are state-varying due to the nonlinearity. Let **x**(*t*) be a temporally evolving variable from some initial state **x**(0), then:

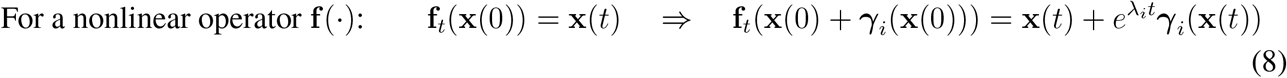

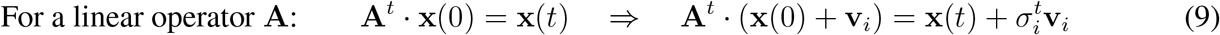

The algorithm to calculate the LEs and CLVs is detailed in ^48, 49^. We described it briefly here. Notice that any random perturbation (δ**x**(0)) would eventually converge to the direction of the fastest growth (***γ***_1_) with growth rate of the leading LE (*λ*_1_). Therefore, the leading LE and CLV can be calculated from numerically simulating the evolution of random perturbations:

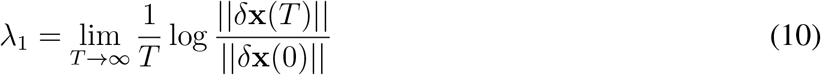

To calculate the rest of the LE spectrum (*λ*_2_, · · ·, *λ*_*N*_), the challenge is that unless the initial perturbation happens to be exactly perpendicular to the direction of the leading CLV, then the evolution of this perturbation would eventually be dominated by the leading direction. In other words, to reveal the growth rate of the second leading direction, we need to remove the effect of the leading direction by perturbing in a subspace orthogonal to the leading direction. In practice, to numerically calculate the first *K* LEs, we simulated the evolution of *K* small random perturbations on a chosen trajectory in parallel. During simulation, we periodically re-orthogonalized the *K* deviation vectors to prevent their collapse onto the fastest-growing direction. For orthogonalization, we did QR factorization in a way that the *ith* deviation vector is projected to the subspace orthogonal to all the 1, · · ·, *i* − 1 vectors. The growth rate of such projection gave the *ith* LE. For each session, we randomly picked *A* = 50 correctly performed trials and used the agent simulations as the unperturbed trajectories. For each trajectory, we repeated the above process with random initial perturbations for *B* = 100 times. In figure 3 we averaged across all sessions, trajectories and repeats:

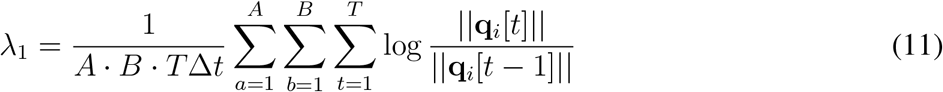

where **q**_*i*_ is the *ith* deviation vector after orthogonalization. However, note that **q**_*i*_’s are no longer covariant with the dynamics due to the orthogonalizations. To reveal the true CLVs, we need to recombine {**q**_*i*_[*t*]} by:

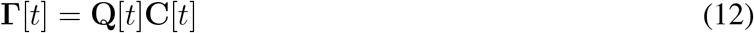

where **Γ**[*t*] contains column vectors of the CLVs (***λ***_*i*_[*t*]) and **Q**[*t*] contains column vectors of **q**_*i*_[*t*]. The coefficient **C**[*t*] is an upper triangular matrix that recombines the orthogonal {**q**_1_[*t*], · · ·, **q**_*i*_[*t*]} to recover the *ith* CLV ***γ***_*i*_[*t*]. This is to counteract the orthogonalizing process during forward simulation. **C**[*t*] is calculated in reverse iterations using **C**[*t*] = **R**^*−*1^[*t*]**C**[*t* + 1]**D**[*t*], where **R**[*t*] is the upper triangular matrix in QR factorization that orthogonalizes **q**_*i*_[*t*]’s during simulation. To ensure that the CLVs have unit norms, **C**[*t*] is normalized column-wise by **D**[*t*], a diagonal matrix containing column norms of **R**^*−*1^[*t*]**C**[*t* + 1].

### Weight analyses

For RNN weight *J*_*ij*_, we call neuron *i* its target neuron and neuron *j* its source neuron. RNN weights were grouped according to the source and target neurons’ peak forward positions and peak lateral velocities. For all panels in Figure 4 f-i except the right plots in panel 4g and 4i, we pooled weights across all agents for individual sessions and all the CV folds. For figure 4f-g, we took a forward position bin of 5 cm, and determined the peak position of neurons based on their trial-averaged activity. For figure 4 h-i, we took a wider forward position bin of 10 cm and used the lateral velocity bin of 0.1, and determined the peak forward position and peak lateral velocity of a neuron as the 2-D indexes of its peak conjunctive activity in bins of forward position and lateral velocity.

### Adding bias to the weights in the data-derived agents

The influence of inhibition strength on neural dynamics was examined for the trial period after cue offset, in order to isolate the constraining effect of visual cues. Therefore, the neural and behavioral trajectories in the cue period were held fixed, and re-training and evaluations were performed following cue offset. We started with the input, recurrent, and output weights of the data-derived agent. A bias (chosen from [−0.02, −0.01, 0.005, 0.01]) was added to the recurrent weights from the time step of cue offset, instantaneously changing the neural activity and behavioral outputs. Then, the same procedure of online imitation learning was performed. During this re-training phase, the input and biased recurrent weights were held fixed and only the velocity output weights were re-trained to minimize the difference between the new velocity outputs (forward and lateral) and those of the original agent. Evaluations of neural effects (Fig. 5j-l) were performed for the arm epoch.

### Toy model with fixed random weights

For the toy model in Figure 5a-f, the within-pool weights were drawn from 𝒩 (*σ*(*β* + *µ*), *σ*^2^) and the across-pool weights were drawn from 𝒩 (*σβ, σ*^2^). Here, the mean was intentionally scaled by *σ*, as the eigen-spectrum properties of Gaussian matrices usually depend on not the absolute mean, but its proportion to the standard deviation. A fixed positive *µ* = 0.3 was considered to impose the competition motifs. To evaluate the dynamical properties of a specific combination of *µ, σ* and *β*, a specific weight matrix realization was drawn from the Gaussian distributions, and multiple trial trajectories were simulated based on the same weight realization from different random activity initializations. The same activation function, time constant and integration time step as in the data-derived agents were used for the toy model. Phase transitions (Fig. 5f, Supplementary Fig. 6) combined the results from 10 random weight realizations for each parameter combination of *σ* and *β*, each simulated for 100 left and 100 right trials, with a constant cue signal presented for 400 time steps. For Lyapunov exponent analyses (Fig 5. d,e), numerical simulation with a given weight realization and constant cue inputs was performed from a single activity initialization for 50000 time steps to provide thorough sampling of the state space and complete convergence of the estimated Lyapunov exponents. Residual PCs were computed based on the last 20000 time steps of the simulation. Reported Lyapunov exponent analyses results combined simulations from 50 random weights realizations.

### Reward-based learning with the Actor-Critic algorithm

For reward-based learning of obstacle avoidance behavior, two obstacles were placed at forward position 180, lateral span [−7, 16], and forward position 230, lateral span[15, 35] in the arm epoch. Note that the arm epoch started at forward position 150 and ended at 235, with allowed range for lateral movement spanning [−35, 35], all units in cm. Because the two obstacles overlapped in their lateral spans and the upper one connected with the upper boundary of the arena, such arrangement of obstacles made sure that they could not be avoided by simply adding a bias to the lateral velocity output.

The input weights, recurrent weights, and output weights to forward velocity of the data-derived agent were held fixed while the output weights to lateral velocity were re-trained using the actor-critic method as in reference ^58^. Here we described the setup briefly. The actor, simply being the lateral velocity output layer, generated lateral velocity *a*_*t*_ from RNN activity **r**_*t*_. The actor weights were initialized as that of the original data-derived agent. The critic was a multi-layer perceptron (MLP) with 2 hidden layers (layer width 64 and 16), and was trained to predict a value *Q*_*t*_ based on the state **s**_*t*_, action **a**_*t*_ and desired choice **c**. Here the state was defined as the forward and lateral positions in the maze, both expanded by a set of 5 cosine basis functions, and the action was the lateral velocity, expanded by a set of 10 cosine basis functions. The choice was represented as a binary variable, [1 0] or [0 1]. These together constituted the input (∈ ℝ^22^) for the critic network. For training, we considered trials simulated from the initial conditions sampled from the first data-points of mouse trials. Trained together, the actor was optimized to maximize the value of its actions, judged by the critic, while the critic was optimized to minimize the reward prediction error. The actor and critic networks were updated once per trial, as they were trained on a replay buffer which contained the entire input and output sequences for both networks at each time point of the trial, as well as the reward history if any.

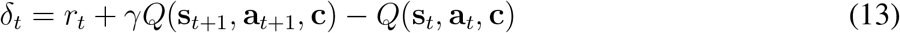

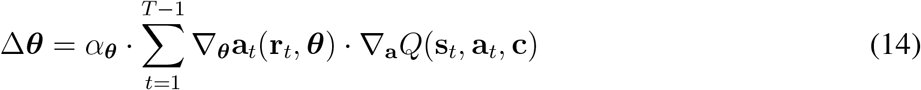

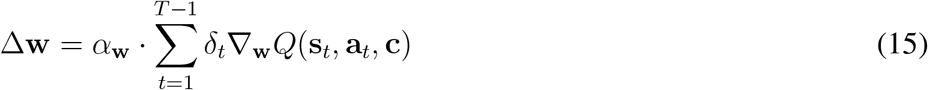

where ***θ*** stands for the parameterization of the actor, and **w** stands for the parameterization of the critic. The reward, *r*_*t*_, might occur at 3 positions along the maze: (1) Reward of getting over the first obstacle, evaluated at 181 cm. Successful avoidance led to a transient reward of *r*^1^. Hitting the obstacle led to game-over and resulted in a negative reward of *r*^neg^. (2) Reward of entering the correct arm evaluated at 220 cm. Correctness led to a transient reward of *r*^2^, while incorrectness led to a transient negative reward of *r*^neg^. (3) Reward of getting over the second obstacle and getting to the correct goal location, evaluated at the end of maze. If success, agent received a reward of *r*^3^. If hitting the second obstacle or choosing the wrong goal, agent received *r*^neg^. We used the Adam optimization method implemented in PyTorch. 5-fold CV was performed with the same ways of train-test splits as in the data-derived agent. Hyper-parameters, including the amplitude of the rewards [*r*^1^, *r*^2^, *r*^3^, *r*^neg^] and the learning rates of the actor and critic networks [*α*_***θ***_, *α*_**w**_], were searched based on the training performance in the first session of each mouse and then generalized to the other sessions. To facilitate learning, we implemented a curriculum learning procedure, that the length of the obstacles were gradually increased to the full length. Agents were trained in each curriculum stage for a maximum of 30 epochs and would be promoted early to the next stage if training performance reached 85%. We also implemented curriculum rollback, that the agents would be reverted to the previous checkpoint if performance stayed low for more than 4 epochs. The best checkpoint for each session was selected based on training performance across left and right trials, and we reported the cross-validated testing performance for left trials.

### Supervised learning in the obstacle maze

RNN activity in the original maze was recorded and fixed. Output weights to the lateral velocity were retrained to minimize the loss in the obstacle maze. The loss was computed in the following way: Assuming the pace of forward progressing was unchanged, use the new lateral velocity to simulate lateral locomotion in the maze. The loss of a trial evaluated whether the trajectory arrived at the right Y-arm and whether the trajectory went through an obstacle, where the later term took the form of softplus(min(*y*^upper^−*y, y*−*y*^lower^)). Here *y*^upper^ and *y*^lower^ are the upper and lower lateral position boundary of an obstacle, and *y* is the lateral position when a trajectory arrived at the obstacle. The term is positive when *y* is between *y*^upper^ and *y*^lower^, which means the trajectory collided with the obstacle. The output weights were trained on both left and right trials for 10 epochs with Adam optimization in Pytorch. We reported the cross-validated testing performance of left trials, averaged across 10 repeats with random seeds.

### Dynamic Mode Decomposition (DMD)

We performed exact DMD ^62^ on the RNN activity of the data-derived agents. In brief, DMD computes the best-fit one-step linear map of the data. Note that DMD does not assume linear dynamics, but rather finds the linear transformation that best captures the dominant, repeatable patterns in nonlinear dynamics. We started by arranging the RNN activity 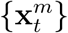 (time step *t* in trial *m*) into data matrices **X** and **Y**: 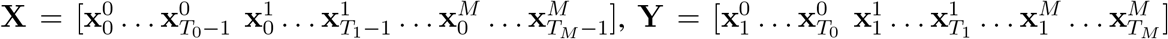, so that **X** and **Y** concatenate all trials with a 1-step offset. We define operator **A** to be the least-squares solution to the problem **AX** = **Y**. Then the DMD modes and eigenvalues of the system are given be the eigenvectors and eigenvalues of **A**. In practice, exact DMD was calculated by the following algorithm from reference ^62^, described briefly here: (1) Compute the reduced SVD of **X**: **X** = **UΣV**^*∗*^ that preserves 99.9% of the energy. (2) Define matrix **Ã** = **U**^*∗*^**YVΣ**^*−*1^. Compute eigen-decomposition of **Ã Ã w** = *λ***w**. Each nonzero *λ* is a DMD eigenvalue. (3) The DMD mode corresponding to *λ* is computed as ***ϕ*** = **YVΣ**^*−*1^**w**. The DMD eigenvalue, *λ*, can be written in polar coordinate *λ* = *ρe*^*itθ*^, where the real part log *ρ* gives the exponential growth rate and the imaginary part *θ* gives the oscillation frequency. One-step DMD prediction error was calculated by 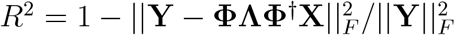, where **Φ** = [***ϕ***_1_ … ***ϕ***_*r*_] and **Λ** = Diag(*λ*_1_ … *λ*_*r*_).

In theory, by tuning the linear readout weight from the RNN, one could combine the oscillatory components with specific loading and phase offset for each component, creating a wide range of output functions. Here we sketch out the main idea. Any initial state can be expressed with the basis of DMD modes, 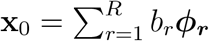. The linear evolution of the activity is then given by 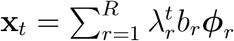, and the linear readout with weight **c** is 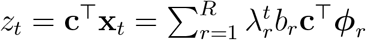. For a pair of complex eigenvalues, *λ* = *ρe*^*itθ*^ with mode ***ϕ*** and amplitude *b*, and its conjugate 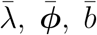, their contribution to the output is given by

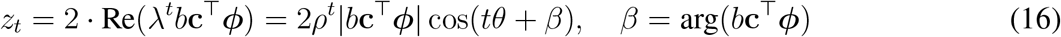

Thus, specific choice of **c** determines *b***c**^⊤^***ϕ***, whose magnitude and argument sets the loading and phase offset of the component associated with ***ϕ***. Therefore, having more slowly decaying components (that we define as −0.2 ≤ *ρ* ≤ 0.2) is beneficial to the system’s capacity to generate arbitrary targets.

## Supplementary Information

### Hyperparameters

|  | Data-derived agents | Toy model |
| --- | --- | --- |
| Weights trainable | Yes | No |
| External inputs | Dependent on RNN outputs | Fixed pattern |
| Number of RNN units | $2628 \pm 506$ (mean ± s.d.) | 400 |
| Number of input channels | 11 | 1 |
| Number of output channels | 3 | 0 |
| Integration time step ($dt$) | 0.0093s | 0.0093s |
| Target time step | 0.186s | N/A |
| Time constant ($\tau$) | 0.1s | 0.1s |
| Learning rate (recurrent weights) | 5 | N/A |
| Learning rate (output weights) | 1 | N/A |
| Noise during training | 0.1 | N/A |
| Additional L2 reg. on input weights | 500 | N/A |

### Removal of RNN self-projections

For this section, we only consider the recurrent weights. In an RNN with *N* units, for each unit, there are only *N* − 1 pre-synaptic units since we remove its self-projection. So the corresponding inverse correlation matrix for post-synaptic unit *i* (without considering itself) should be:

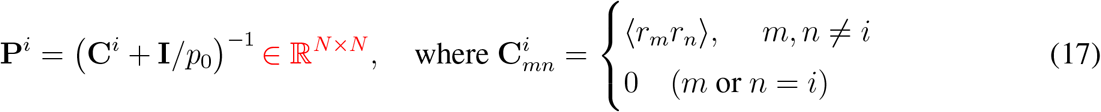

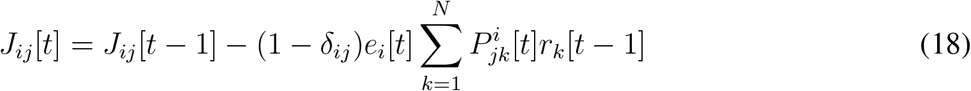

Note that although **C**^*i*^ can be constructed from the sub-matrix of **C, P**^*i*^ can NOT be constructed from the sub-matrix of **P**.

#### Method 1: Directly update individual **P**^*i*^ ‘s

For notation simplicity we define

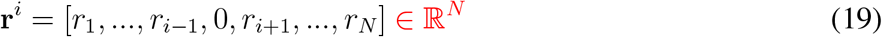

Then the update rule should be

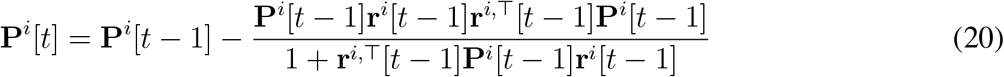

Limitation: This method requires maintaining all the 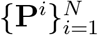 in the memory. When *N* is large (~ 3000 in our case), this would require ~ 200 GB memory.

#### Method 2: Derive {**P**^*i*^} from the original **P**

Another idea is to do a bit more computation in exchange for less required memory.

For notation simplicity, define 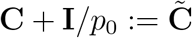. Then

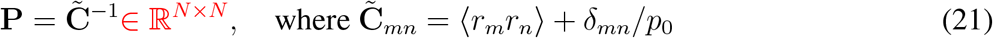

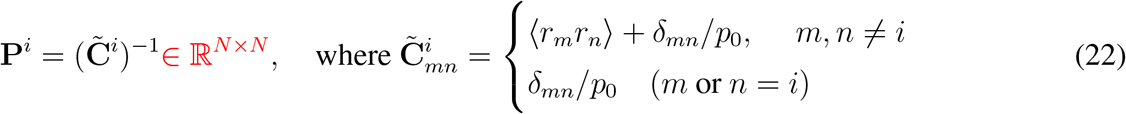

Note that 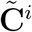 can be constructed from the sub-matrix of 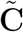:

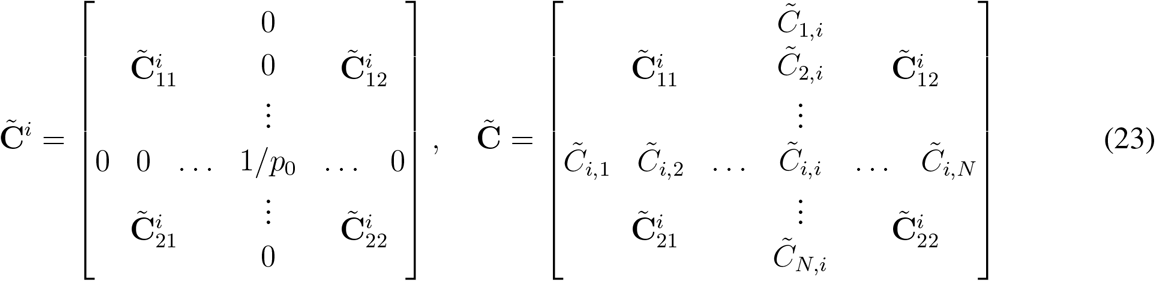

Note that 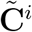 differs from 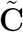by a rank-two operation:

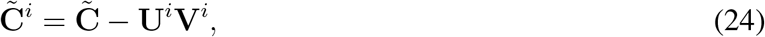

where

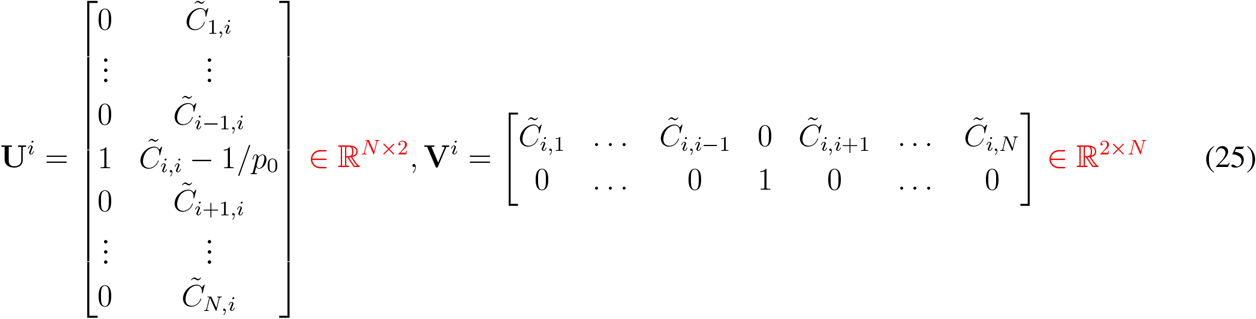

Using the Woodbury formula, 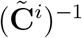 can be derived from 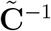:

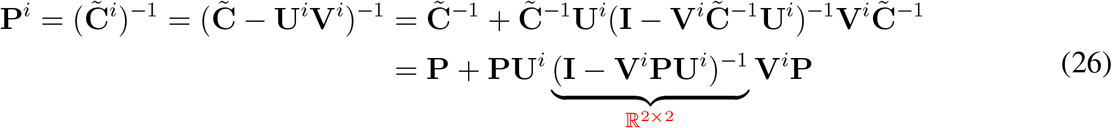

The advantage of this method is that all the 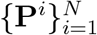 are temporary variables and do not need to constantly occupy the memory.

We can use a compact matrix formula to speed up the computation, using ⊙ to denote element-wise product (broadcast may apply) and ×1 in the matrix shape to denote the operation of adding a dimension. Say there are *M* post-synaptic units in total.

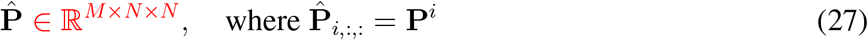

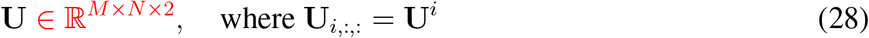

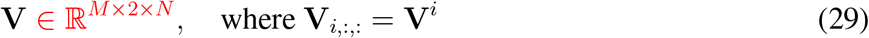

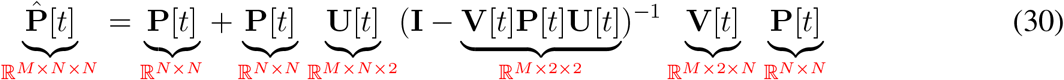

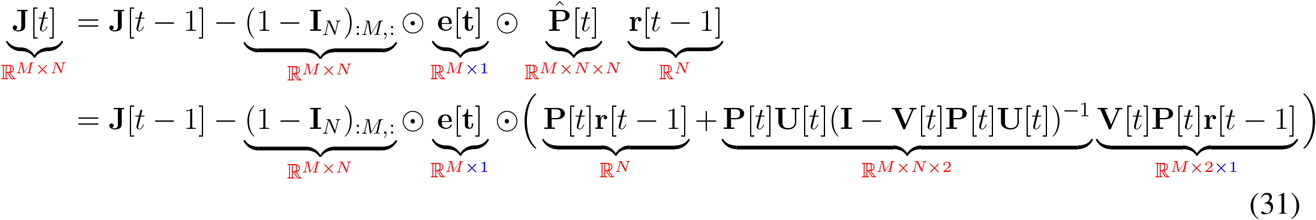

As indicated by the under-braces, if one performs the matrix multiplications in a proper order, we can avoid directly computing the big 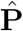matrix, and there is no need to save it in the memory even temporarily! Therefore the memory/space required is not much different from the normal training algorithm.

Furthermore, **PU** and **VPU** can be hand-calculated to speed up the program (as they are the only two 𝒪(*MN*^2^) heavy calculations involved):

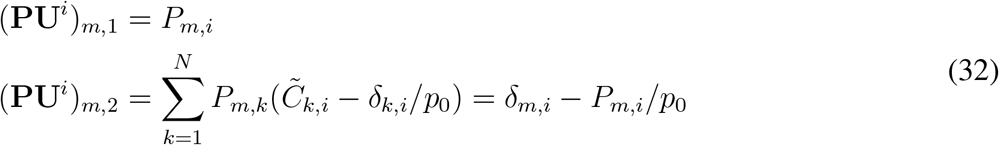

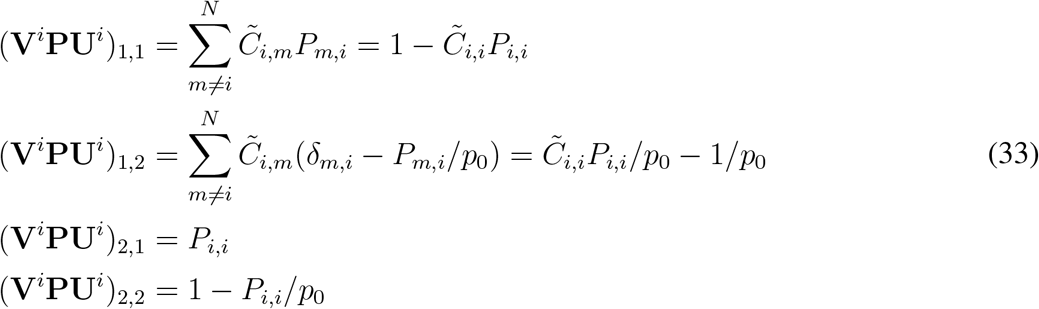

